# Early Embryonic Establishment of Constitutive Heterochromatin Involves H3K14ac-mediated Recruitment of Eggless/SetDB1

**DOI:** 10.1101/2023.12.10.570967

**Authors:** Ruijun Tang, Mengqi Zhou, Yuwei Chen, Zhenghui Jiang, Xunan Fan, Jingheng Zhang, Aiping Dong, Lu Lv, Song Mao, Fang Chen, Jinrong Min, Ke Liu, Kai Yuan

**Affiliations:** Hunan Key Laboratory of Molecular Precision Medicine, Department of Neurosurgery, Xiangya Hospital & Center for Medical Genetics, School of Life Sciences, Central South University, Changsha, Hunan, China; Hubei Key Laboratory of Genetic Regulation and Integrative Biology, School of Life Sciences, Central China Normal University, Wuhan, Hubei, China; Yichun Maternal and Child Health Care Hospital, Yichun, Jiangxi, China; Structural Genomics Consortium, University of Toronto, Toronto, Ontario, China; National Clinical Research Center for Geriatric Disorders, Xiangya Hospital, Central South University, Changsha, Hunan, China; Furong Laboratory, Hunan, China; The Biobank of Xiangya Hospital, Central South University, Changsha, Hunan, China

## Abstract

Constitutive heterochromatin, built on various types of repetitive DNA, is a fundamental feature of eukaryotic nucleus essential for transposon silencing and genome stability. However, the molecular programs driving its *de novo* establishment during early embryogenesis remain poorly understood. Here, we show that histone H3 lysine 14 acetylation (H3K14ac) is maternally inherited and partially colocalizes with hallmarks of constitutive heterochromatin, H3 lysine 9 trimethylation (H3K9me3) and its methyltransferase Eggless/SetDB1, around the mid-blastula transition in *Drosophila* early embryos. Concealing H3K14ac by either antibody injection or maternal knockdown of Gcn5 diminishes Eggless/SetDB1 nuclear foci and reduces the deposition of H3K9me3. Structural analysis reveals that Eggless/SetDB1 recognizes H3K14ac via its tandem Tudor domains, and disrupting the binding interface causes defects in Eggless/SetDB1 distribution and derepression of a subset of transposons. Therefore, H3K14ac, a histone modification associated with active transcription, is a crucial piece of the maternal programs that introduce constitutive heterochromatic features to the newly formed zygotic genome.

## INTRODUCTION

After fertilization, profound epigenetic resetting transforms the genome inherited from two fully differentiated gametes into a relatively naïve state, preparing a blank scroll for the painting of a new life cycle. Basic epigenetic landscapes are then laid out in subsequent embryogenesis following stereotyped developmental programs, including the stepwise package of various repetitive DNA elements into transcriptionally inert constitutive heterochromatin (Allshire and Madhani, 2018; Campos et al., 2014; Liu et al., 2020; Padeken et al., 2022). In *Drosophila melanogaster*, approximately 20% of the genome, composed of large tandem repeats and dispersed transposable elements, initiates heterochromatinization when embryos slow down the cell cycles at the major embryonic transformation period called the mid-blastula transition (MBT) (Armstrong and Duronio, 2019; Seller et al., 2019; Yuan et al., 2016). This establishment of constitutive heterochromatin is orchestrated largely by maternal programs that are of significant complexity (Yuan and O’Farrell, 2016), with Eggless/SetDB1-catalyzed deposition of H3K9me3 being the major chromatin silencing effector (Armstrong and Duronio, 2019; Seller *et al*., 2019). Different repetitive DNA elements seem to take on distinct molecular pathways to arrive at a heterochromatic state. Tandem AAGAG repeat relies on specific recognition by the pioneer factor GAF to initiate H3K9me3 deposition and transcriptional silencing (Gaskill et al., 2023). Some transposable elements, however, utilize piRNA-directed mechanism to engage Piwi and the PICTS complex with the nascent transposon transcripts to recruit Eggless/SetDB1 and instruct local heterochromatin formation in the early embryos (Batki et al., 2019; Fabry et al., 2021; Mugat et al., 2020; Ninova et al., 2020b; Zhao et al., 2019). Besides, the emergence of heterochromatic features on different repetitive DNA elements is heterochronic. A group of tandem repetitive elements such as the 359-bp and AAGAG repeats attain H3K9me3 decoration and HP1a accumulation at the MBT, preceding other repeats (Gaskill *et al*., 2023; Yuan and O’Farrell, 2016). To date, our understanding of the molecular basis underlying the precise recruitment of Eggless/SetDB1 and hence the inheritance of constitutive heterochromatic states across generations is far from complete.

Unlike histone methylation that can decorate active or silent chromatins, acetylation of histone tails is thought to be invariably linked to transcriptional activation (Filion et al., 2010; Jenuwein and Allis, 2001; Kharchenko et al., 2011). For instance, intergenerationally maintained H4K16ac is instructive to the zygotic genome activation during *Drosophila* early embryogenesis (Samata et al., 2020). H3K14 is acetylated mainly by histone acetyltransferase Gcn5, which serves as the catalytic subunit of four different transcriptional coactivator complexes that stimulate gene expression in *Drosophila* (Torres-Zelada and Weake, 2021). Null *Gcn5* alleles block the oocyte development and larva-to-adult metamorphosis (Carre et al., 2005). Transgenic flies carrying histone gene unites (His-GUs) in which H3K14 is substituted for alanine or arginine (H3K14A or H3K14R) in a histone deficiency mutant background manifest lethality at the embryonic or second-instar larval stage (Regadas et al., 2021; Zhang et al., 2019). In late embryonic development, H3K14ac enriches over the gene body of a group of tissue-specific genes, defining a unique chromatin state essential for organogenesis (Regadas *et al*., 2021). Intriguingly, *in vitro* peptide binding assay revealed that human SetDB1, a methyltransferase of H3K9me3, can interact with doubly modified H3 tails containing H3K14ac and H3K9me2/3 through its N-terminal tandem Tudor domains (TD). Crystal structure demonstrated that the interface between TD2 and TD3 of SetDB1 forms the binding pockets for these modifications (Jurkowska et al., 2017). These observations suggest a functional connection between H3K14ac and the repressive H3K9me3 histone mark. However, the early embryonic dynamics of H3K14ac relative to H3K9me3, as well as the potential biological significance of H3K14ac in the establishment of constitutive heterochromatin, remain undetermined.

Here, we show that in *Drosophila* H3K14ac is transmitted from the female germline to the zygotic embryos, where it is recognized by the tandem Tudor domains of Eggless/SetDB1, facilitating the concentration of Eggless/SetDB1 on repetitive DNA elements. Either erasing the H3K14ac modification or disrupting the recognition of H3K14ac by Eggless/SetDB1 leads to reduced deposition of H3K9me3 and derepression of a group of transposons, compromising the integrity of constitutive heterochromatin and organismal viability. Our study identifies the H3K14ac-mediated recruitment of Eggless/SetDB1 as an important module of the molecular machinery underneath the intergenerational inheritance of the constitutive heterochromatic states.

## RESULTS

### H3K14ac is maternally inherited and colocalizes with H3K9me2/3 and Eggless/SetDB1 at the apical pole of nuclei around the mid-blastula transition

*Drosophila* early embryogenesis begins with rapid nuclear divisions, deferring the emergence of constitutive heterochromatic hallmarks in the zygotic genome until the cell cycles slow down at the MBT (Seller *et al*., 2019; Yuan and O’Farrell, 2016). We set out to characterize the spatial distribution of H3K14ac in the syncytial and cellular blastoderm embryos in comparison to H3K9me2/3 (Figure 1A). While H3K9me2 histone modification became detectable slightly earlier than H3K9me3, both repressive marks progressively accumulated in cycle 13 and cycle 14 embryos at the apical pole of nuclei where numerous repetitive DNA elements reside. H3K14ac was already abundant in nuclei of early syncytial blastoderm embryos (cycle 11-12), forming several nuclear puncta in addition to the ubiquitous nuclear distribution. In cycle 13 and cycle 14 embryos, H3K14ac became enriched at the apical regions of nuclei, colocalizing with the H3K9me2/3 modifications (Figure 1A). We quantified the H3K14ac antibody staining signals within and without the H3K9me3-decorated compartments in cycle 14 embryos, and detected that the mean fluorescent intensity of H3K14ac was significantly higher in the H3K9me3-positive regions (Figure 1B). Eggless/SetDB1 is the major histone methyltransferase in the early embryos to deposit H3K9me2/3 modifications (Seller *et al*., 2019). We collected embryos from a knock-in *Drosophila* line in which the Eggless/SetDB1 is tagged with GFP at its N terminus (GFP-SetDB1), and performed immunostaining to visualize H3K14ac. Similar to H3K9me3, GFP-SetDB1 accumulated at the apical poles of nuclei in cycle 14 embryos and significantly overlapped with H3K14ac (Figure 1A and 1C).

**Figure 1.**
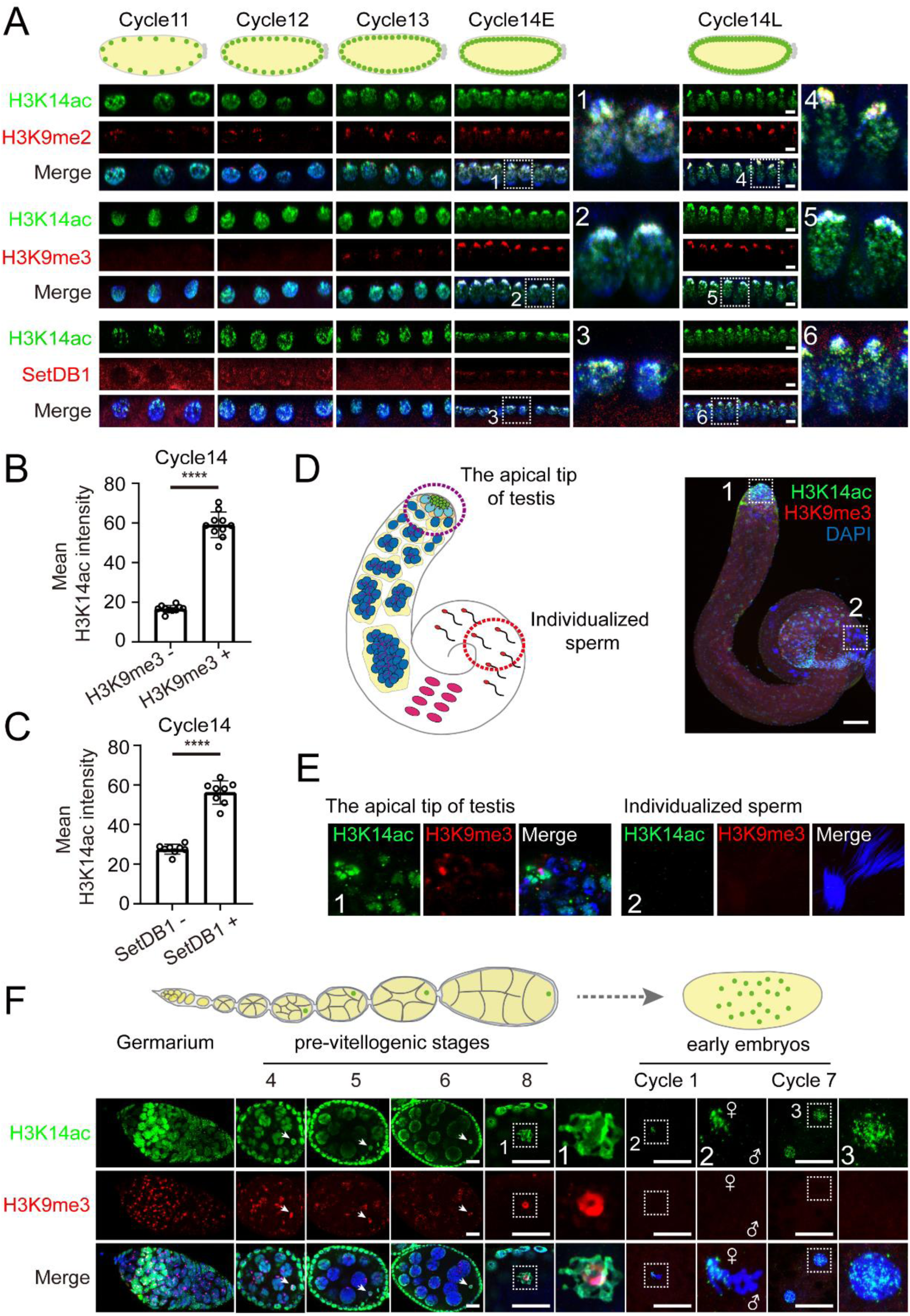
Distribution of H3K14ac during *Drosophila* gametogenesis and early embryogenesis. **(A)** Immunofluorescence of fixed embryos at different developmental stages. The estimated embryonic cell cycles, determined using either internuclear distances for the syncytial blastoderm embryos or nuclear lengths for the cellular blastoderm embryos, are labeled at the top. The antibody staining of H3K14ac is shown in green, and H3K9me2/3 or SetDB1 in red. DAPI-stained DNA is shown in blue. The numbered images show enlarged views of the corresponding dashed line-labeled regions. Bars: 5 μm. **(B)** Quantification of nuclear H3K14ac fluorescent signals in H3K9me3 positive (H3K9me3 +) and H3K9me3 negative (H3K9me3 -) regions in cycle 14 embryos. Unpaired *t*-test, ****: *p* < 0.0001. Error bars represent the SD. **(C)** Quantification of nuclear H3K14ac signals in SetDB1 positive (SetDB1 +) and negative (SetDB1 -) regions in cycle 14 embryos. Unpaired *t*-test, ****: *p* < 0.0001. Error bars represent the SD. **(D)** Immunofluorescence of *Drosophila* testis with H3K14ac (green) and H3K9me3 (red) antibodies. DNA was labeled with DAPI (blue). A cartoon of *Drosophila* testis is shown on the left. The dashed line-labeled regions are zoomed-in in (E). Bar: 20 μm. **(E)** H3K14ac and H3K9me3 fluorescent signals at the apical tip of testis and in individualized sperms. **(F)** Immunofluorescence of *Drosophila* ovaries and early embryos with H3K14ac (green) and H3K9me3 (red) antibodies. DNA was stained with DAPI (blue). The numbered images show enlarged views of the dashed line-labeled regions. Metaphase chromosomes of paternal or maternal origin in cycle 1 embryos are labeled with ♂ or ♀ respectively. Bars: 15 μm.

Since H3K14ac was already detectable in cycle 11 embryos, we traced its presence during gametogenesis as well as in pre-blastoderm embryos. *Drosophila* testis is a blind-ended coiled tube, with the germline stem cells at the apical tip and matured sperms in the distal regions (Figure 1D). While both H3K14ac and H3K9me3 were detected in cells at the testis tip, these two histone modifications showed little overlap. However, neither of these modifications was maintained in the individualized sperms (Figure 1E). *Drosophila* ovary is composed of a bundle of ovarioles in which the stem cells are located at the tip of the germarium and the developing egg chambers are arranged in a linear manner. The oocyte localizes to the posterior side of each egg chamber (Figure 1F). We detected strong H3K14ac and H3K9me3 signals in oocytes at different developmental stages. The H3K9me3 was concentrated in the middle of the germinal vesicle, whereas H3K14ac appeared more expanded and amorphous in its distribution (Figure 1F, pre-vitellogenic stage 8). Therefore, both H3K14ac and H3K9me3 were absent in mature sperms but present in oocytes. After fertilization, the chromosomes of maternal and paternal origins are still kept separate in mitosis of the first cell cycle. Compared to H3K9me3, which was removed from all the zygotic genome, the H3K14ac signal was detected on half of the metaphase chromosomes that were likely inherited from the maternal pronucleus. Subsequently, the H3K14ac decorated the zygotic nuclei throughout the pre-blastoderm stage (Figure 1F). These results indicate that H3K14ac is maternally transmitted, persists throughout the early embryogenesis, and partially colocalizes with hallmarks of constitutive heterochromatin during the MBT time frame.

### Masking or erasing the H3K14ac histone code disrupts Eggless/SetDB1 nuclear foci formation in syncytial blastoderm embryos

The colocalization of H3K14ac with Eggless/SetDB1 and H3K9me3 during the period of heterochromatin formation prompted us to investigate its functional significance. To this end, we injected H3K14ac or H3K9me3 antibody into embryos at the mitosis of cycle 11 to mask the modification, and monitored the localization of the endogenously tagged GFP-SetDB1 as well as RFP-HP1a expressed from a transgene (Figure 2A). Compared to the control, injection of H3K9me3 antibody showed little effect on the nuclear foci formation of GFP-SetDB1 and RFP-HP1a until the embryos developed to the interphase of cycle 13, during which the presence of H3K9me3 antibody mildly reduced the number of GFP-SetDB1 as well as RFP-HP1a foci (Figure 2B, Supplementary Movies S1-S2). Notably, injection of the H3K14ac antibody gave rise to a stronger phenotype. The nuclear foci of GFP-SetDB1 and RFP-HP1a were significantly disrupted in the interphases of both cycle 12 and cycle 13 embryos, and the chromosome segregations in mitosis 12 were unsuccessful (Figure 2B-2C, Supplementary Movie S3).

**Figure 2.**
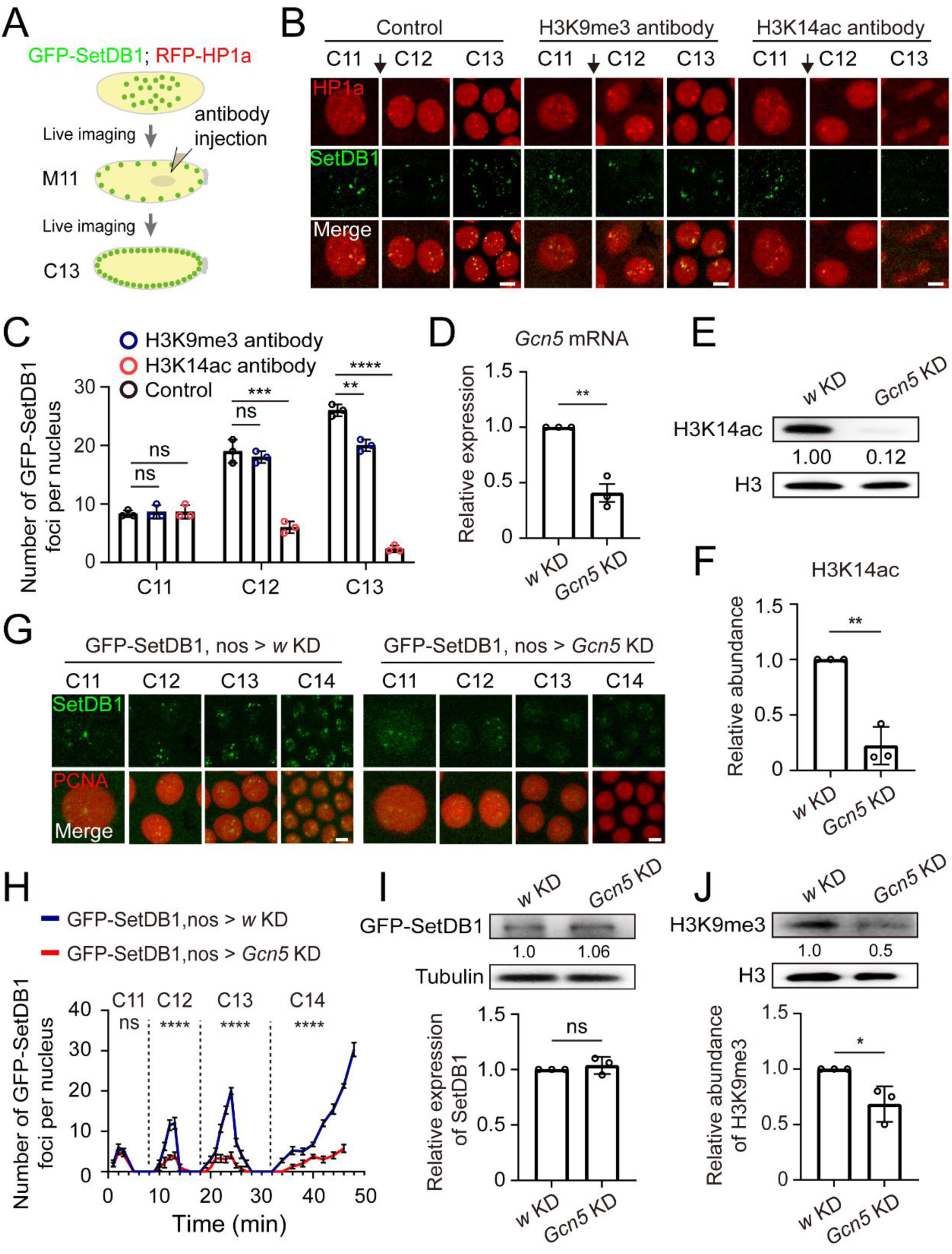
Perturbation of H3K14ac impacts SetDB1 localization during early embryonic development. **(A)** Schematics of live imaging experiments with antibody-injected embryos. **(B)** Time-lapse images of nuclei in embryos expressing mRFP-HP1 (red) and GFP-SetDB1 (green). H3K9me3 or H3K14ac antibody diluted in PBS was injected during mitosis 11 (black arrows). PBS was served as a control. Bars: 5 μm. **(C)** Quantification of the numbers of nuclear GFP-SetDB1 foci in the embryos before and after injection. Unpaired *t*-test, ns: no significance, **: *p* < 0.01, ***: *p* < 0.001, ****: *p* < 0.0001. Error bars represent the SD. **(D)** RT-qPCR analysis of *Gcn5* mRNA abundance in 0-2 h embryos from *GFP-SetDB1*, *nos-Gal4* > *w* (*w* KD) or *GFP-SetDB1*, *nos-Gal4* > *Gcn5* (*Gcn5* KD) parents. Unpaired *t*-test, **: *p* < 0.01. Error bar represents the SD. **(E)** Western blot analysis of *w* KD and *Gcn5* KD 0-2 h embryos with H3 and H3K14ac antibodies. **(F)** Quantification of the western blot results. Unpaired *t*-test, **: *p* < 0.01. Error bar represents the SD. **(G)** Time-lapse images of nuclei in embryos harboring GFP-SetDB1 (green) and mCherry-PCNA (red) after *w* or *Gcn5* knockdown. Bars: 5 μm. **(H)** Line charts represent the numbers of nuclear GFP-SetDB1 foci in *w* KD and *Gcn5* KD embryos from embryonic cycle 11 to cycle 14. Unpaired *t*-test, ns: no significance, ****: *p* < 0.0001. Error bars represent the SD. **(I)** Western blot analysis of *w* KD and *Gcn5* KD 0-2 h embryos with GFP antibody. The protein level of the endogenously-tagged GFP-SetDB1 shows no statistical difference. Unpaired *t*-test, ns: no significance. Error bar represents the SD. **(J)** Western blot analysis of *w* KD and *Gcn5* KD 0-2 h embryos with H3 and H3K9me3 antibodies. The H3K9me3 modification is downregulated in *Gcn5* KD embryos. Unpaired *t*- test, *: *p* < 0.05. Error bar represents the SD.

To further ascertain the contribution of H3K14ac to the nuclear foci formation of Eggless/SetDB1 and HP1a, we knocked down the expression of the major H3K14ac acetyltransferase Gcn5 in the ovaries and early embryos using *nos-Gal4* (Figure 2D). In these embryos, the level of H3K14ac modification was decreased (Figure 2E-2F), and the accumulation of GFP-SetDB1 on nuclear foci in interphases of cycle 12 to cycle 14 was significantly diminished (Figure 2G-2H, Supplementary Movies S4-S5). While the protein level of GFP-SetDB1 remained unaffected by maternal knockdown of Gcn5 (Figure 2I), the H3K9me3 deposition was reduced, likely attributed to the compromised nuclear foci formation of Eggless/SetDB1 (Figure 2J).

### Erasing the H3K14ac by Gcn5 knockdown compromises transposon silencing and heterochromatin integrity

The disruption of Eggless/SetDB1 localization and reduction of H3K9me3 after maternal Gcn5 knockdown indicated errors in the establishment of heterochromatic silencing. We collected control and Gcn5 knockdown embryos at cycle 14 or embryonic stage 5, and performed RNA-sequencing (RNA-seq) analysis to examine the expression of repetitive DNA elements. A total of 3127 repeat loci exhibited an upregulated expression level in Gcn5 knockdown embryos (Fold Change ≥ 1.5, *p* < 0.05), among which 1067 loci were satellite elements and transposons belonging to 122 different classes (Figure 3A, Supplementary Table S1). For validation, we selected several affected transposons, including *gypsy*, *blood*, *BURDOCK*, and *mdg1*, and confirmed their increased transcription by RT-qPCR in embryos with Gcn5 knockdown (Figure 3B).

**Figure 3.**
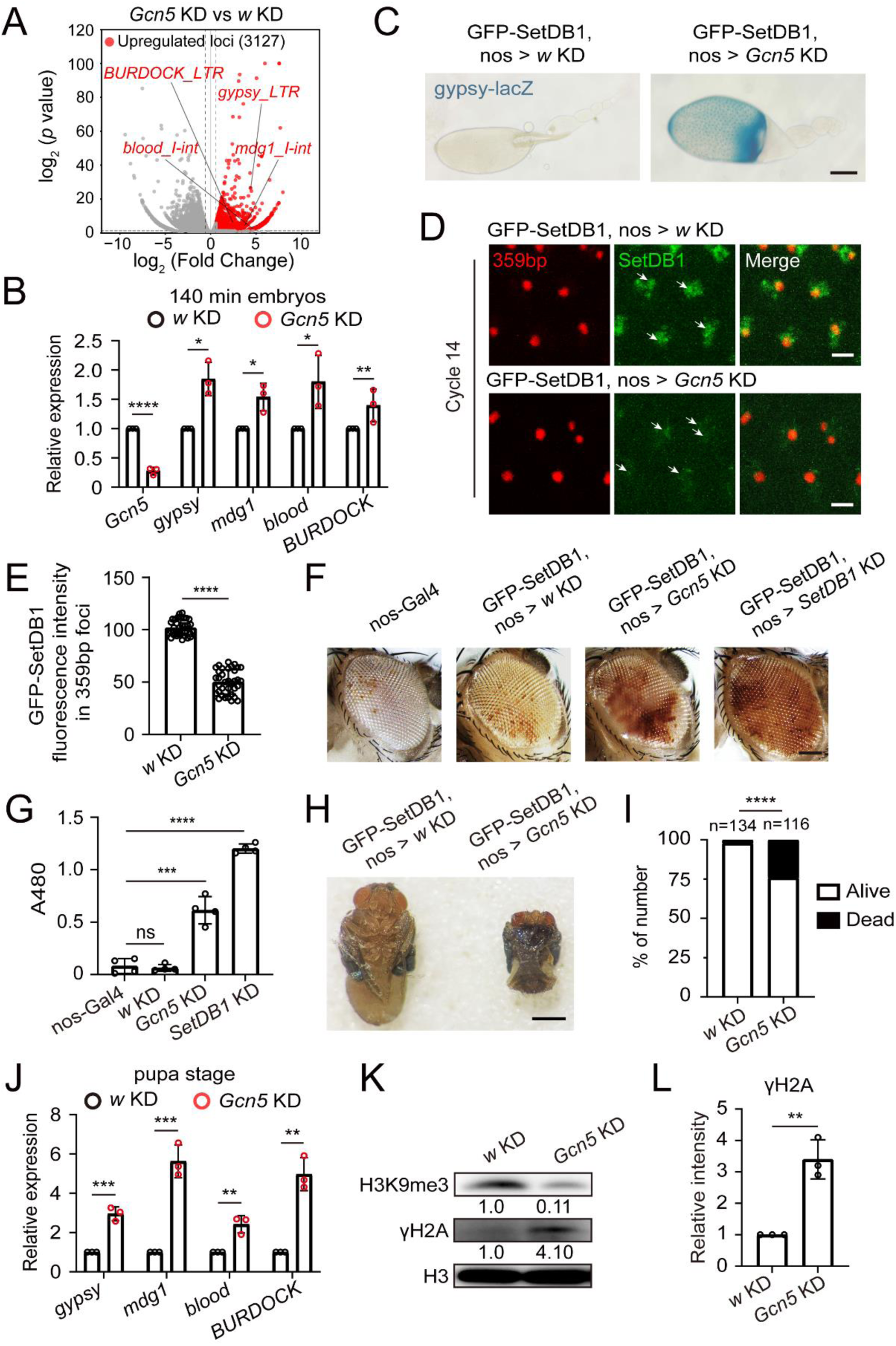
*nos-Gal4* driven *Gcn5* knockdown compromises constitutive heterochromatin. **(A)** Volcano plot of repeats expressions in *w* KD control versus *Gcn5* KD embryos (stage 5, 140 min). Upregulated repeat loci after *Gcn5* KD (Fold Change ≥ 1.5, *p* < 0.05) are plotted in red. **(B)** RT-qPCR validation of upregulation of 4 representative transposons (*gypsy*, *mdg1*, *blood*, and *BURDOCK*) in *Gcn5* KD embryos. Unpaired *t*-test, *: *p* < 0.05, **: *p* < 0.01, ****: *p* < 0.0001. Error bars represent the SD. **(C)** Derepression of *gypsy* in the ovaries after *Gcn5* KD. X-gal staining was used to detect the expression of β-galactosidase from the *gypsy*-*LacZ* reporter (indigo). Bar: 50 μm. **(D)** Video frames from live imaging experiments showing reduced recruitment of GFP-SetDB1 (green) to the 359-bp satellite sequences in *Gcn5* KD embryos. The 359-bp loci were visualized by its TALE-light probe (red). The white arrows indicate the 359-bp regions. Bars: 5 μm. **(E)** Quantification of GFP-SetDB1 fluorescent signals within 359-bp regions in *w* KD and *Gcn5* KD embryos. Unpaired *t*-test, ****: *p* < 0.0001. Error bars represent the SD. **(F)** Eye pictures from female *w^m4h^* (inversion on chromosome X) PEV reporter flies with *nos-Gal4* driven knockdown of *w*, *Gcn5*, or *SetDB1* respectively. Bar: 50 μm. **(G)** Pigment assay of *w^m4h^* reporter flies after knockdown of the indicated genes. Pigment was extracted and the OD at 480 nm was measured. Unpaired *t*-test, ns: no significance, ***: *p* < 0.001, ****: *p* < 0.0001. Error bars represent the SD. **(H)** Impaired pupa development in *nos-Gal4* driven *Gcn5* KD flies. Bar: 100 μm. **(I)** Quantification of successful eclosions in *w* KD control versus *Gcn5* KD flies. Chi-square test, ****: *p* < 0.0001. **(J)** RT-qPCR analysis of transposons expression in *nos-Gal4* driven *w* KD or *Gcn5* KD flies at pupa stage. Unpaired *t*-test, **: *p* < 0.01, ***: *p* < 0.001. Error bars represent the SD. **(K)** Western blot analysis of pupae from *nos-Gal4* driven *w* KD or *Gcn5* KD flies with H3, H3K9me3, and γH2A antibodies. **(L)** Quantification of γH2A western blot signals of *w* KD and *Gcn5* KD pupae. Unpaired *t*-test, **: *p* < 0.01. Error bar represents the SD.

We further enlisted different reporting systems to evaluate the integrity of transcriptional silencing and heterochromatin on various repetitive sequences at different developmental stages. The abnormal activation of *gypsy* can be detected early during oogenesis by the *gypsy-LacZ* reporter (Sarot et al., 2004). Following Gcn5 knockdown, we observed mild X-gal staining signals in the ovaries, confirming that Gcn5-catalyzed deposition of H3K14ac contributed to the repression of *gypsy* (Figure 3C). Previous TALE-light imaging of tandem satellite repeats revealed that the 359-bp repetitive element recruits Eggless/SetDB1 and starts heterochromatinization during the late syncytial blastoderm stage (Seller *et al*., 2019; Yuan and O’Farrell, 2016). Consistently, significant enrichment of GFP-SetDB1 within the 359-bp repeats region was detected in the interphase of cycle 14 control embryos, however, in embryos with Gcn5 knockdown, the accumulation of GFP-SetDB1 at the 359-bp loci was markedly abrogated (Figure 3D-3E, Supplementary Movies S6-S7). The position-effect variegation (PEV) strain *w^m4h^*, which carries an inversion on the X chromosome that positions the *white* gene close to pericentromeric heterochromatin, is a strongly variegating reporter used to assess heterochromatin integrity (Elgin and Reuter, 2013). Compromised heterochromatic silencing leads to derepression of *white* that causes eye pigmentation (Figure 3F). Maternal knockdown of Eggless/SetDB1 driven by *nos-Gal4* resulted in increased pigment level compared to the control group. A similar but milder increase of pigmentation was also observed in flies with maternal knockdown of Gcn5 (Figure 3G), suggesting that proper Gcn5 activity in oocytes and early embryos is required for ensuring the fidelity of heterochromatin formation.

The compromised heterochromatin integrity after Gcn5 knockdown was associated with decreased viability. Approximately 25% of the progeny from the maternal Gcn5 knockdown flies died at the pupal stage (Figure 3H-3I). We collected these pupae and assessed transposon activation and genome stability. In the offspring pupae from the *nos-Gal4*-driven Gcn5 knockdown parents, elevated transcription of transposons alongside increased DNA damage indicated by γH2A abundance was observed (Figure 3J-3L).

### Gcn5 knockdown results in decreased Eggless/SetDB1 and H3K9me3 on a fraction of transposons and satellites

The establishment of heterochromatic silencing on different portions of the repetitive genome involves diverse molecular programs (Yuan and O’Farrell, 2016). RNA-seq analysis revealed that 1067 transposons and satellite loci manifested elevated transcription upon maternal knockdown of Gcn5 (Figure 3A), suggesting that these repetitive elements are regulated by Gcn5-dependent heterochromatin formation. To elucidate the developmental distribution of H3K14ac as well as H3K9me3 in these genomic regions, we conducted chromatin immunoprecipitation sequencing (ChIP-seq) using hand-staged embryos (Figure 4A), and compared the results with published ChIP-seq datasets of other histone modifications generated with *Drosophila* early embryos (Li et al., 2014).

**Figure 4.**
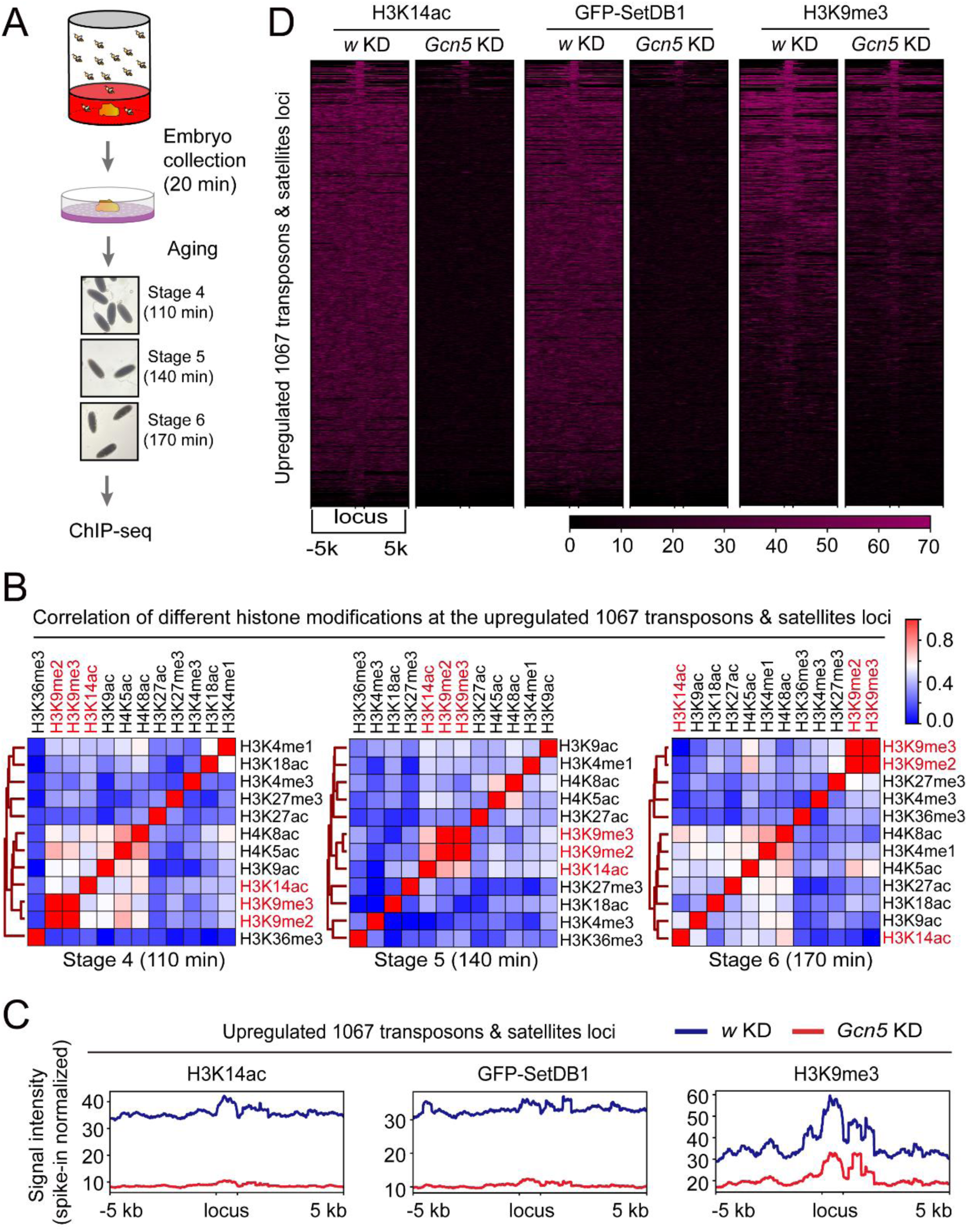
*nos-Gal4* driven *Gcn5* KD results in decreased SetDB1 and H3K9me3 on a fraction of transposons and satellites loci. **(A)** Schematics of embryo collection and preparation for ChIP-seq analysis. **(B)** Heatmaps demonstrating the correlation coefficient among different histone modifications at different embryonic stages on the 1067 upregulated transposons and satellites elements identified in the RNA-seq analysis of *Gcn5* KD and *w* KD embryos. **(C)** Averaged ChIP-seq signals of H3K14ac, GFP-SetDB1, and H3K9me3 around the 1067 upregulated repeats loci in *Gcn5* KD and *w* KD embryos. **(D)** Heatmaps of H3K14ac, GFP-SetDB1, and H3K9me3 ChIP-seq signals around the 1067 upregulated transposons and satellites loci in *Gcn5* KD and *w* KD embryos.

While the genomic distribution of H3K14ac was rather broad, spearman’s correlation analysis revealed that H3K14ac was clustered with H3K9me2 and H3K9me3 as well as several other histone acetylation modifications at the 1067 transposons and satellites loci in stage 4 or syncytial blastoderm embryos (Figure 4B). As embryos developed to stage 5, the critical time window for constitutive heterochromatin formation, the correlation among H3K14ac and H3K9me2/3 became stronger and was exclusive to these three modifications. In stage 6 embryos, H3K14ac was no longer clustered with H3K9 methylation modifications. This dynamic co-occupancy between H3K14ac and H3K9me2/3 at different developmental stages was reminiscent of antibody staining patterns of these modifications in *Drosophila* early embryos (Figure 1A). To directly investigate the impact of H3K14ac on the genomic distribution of Eggless/SetDB1 and H3K9me3, we collected stage 5 embryos from flies with maternal knockdown of Gcn5 and performed ChIP-seq analysis. Gcn5 knockdown resulted in a dramatic decline of H3K14ac genome-wide as well as at the 1067 repetitive loci. Remarkably, GFP-SetDB1 and H3K9me3 were concomitantly downregulated at these genomic regions (Figure 4C-4D).

Together, these results unravel that the Gcn5-catalyzed H3K14ac histone mark is instrumental for the recruitment of Eggless/SetDB1, deposition of H3K9me3, and hence the establishment of heterochromatic silencing on a fraction of repetitive elements during *Drosophila* early embryogenesis.

### Specific recognition of H3K14ac by the tandem Tudor domains of Eggless/SetDB1

We next wanted to unravel the molecular basis underlying this contribution of H3K14ac to the establishment of constitutive heterochromatin in *Drosophila* early embryos. It has been reported that human SetDB1 can recognize the H3K14ac and H3K9me2/3 doubly modified H3 tails *in vitro* via its N-terminal triple Tudor domains (3TD) (Jurkowska *et al*., 2017). To investigate whether *Drosophila* Eggless/SetDB1 could bind H3K14ac, we first conducted an analysis of sequence similarity of the 3TD among *Drosophila*, human, and mouse (Figure 5A). The sequence alignment results revealed that, while the 3TD of human SetDB1 (hSETDB1) exhibited a 99% sequence identity with mouse SetDB1 (mSETDB1), it only displayed a 33% sequence identity with that of *Drosophila* Eggless/SetDB1 (DmSetDB1). Nonetheless, what stood out was the high degree of conservation of amino acid residues responsible for mediating the interaction with H3K14ac within the 3TD across *Drosophila*, mouse, and human (Figure 5A). Therefore, we hypothesized that *Drosophila* Eggless/SetDB1 had the potential to recognize the H3K14ac histone mark through its Tudor domains.

**Figure 5.**
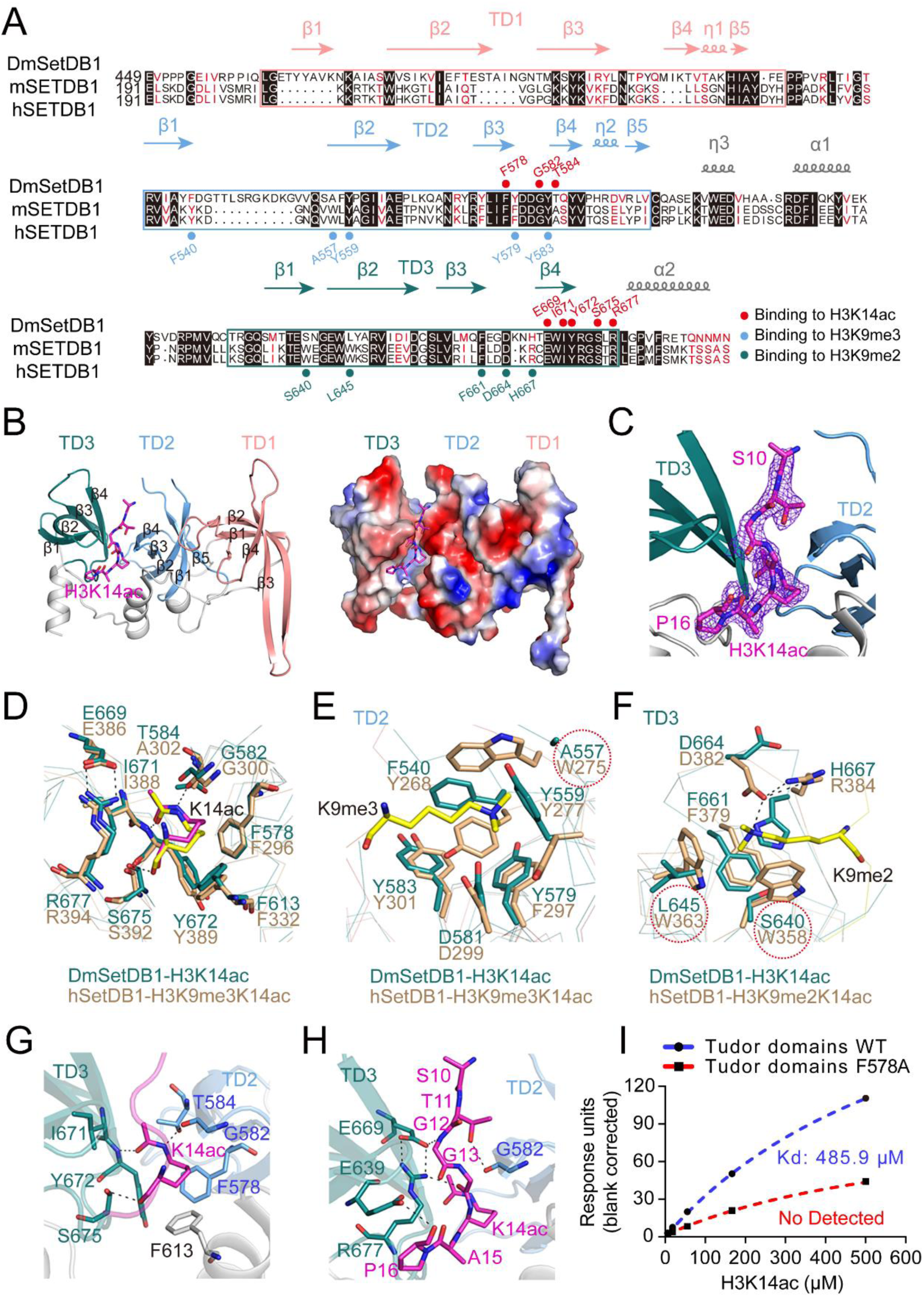
*Drosophila* SetDB1 recognizes H3K14ac modification via its interface between TD2 and TD3. **(A)** Sequence alignment of 3 Tudor domains (3TD) from DmSetDB1, mSETDB1, and hSETDB1. Secondary structure elements of DmSetDB1 3TD are indicated at the top of the sequences. TD1, TD2, and TD3 are colored indian red, cornflower blue, and sea green, respectively. The H3K14ac binding residues of DmSetDB1 are marked by red dots at the top of the sequences. The residues contributing to the H3K9me3 recognition by TD2 of hSETDB1 and their corresponding residues on DmSetDB1 are labeled with cornflower blue dots. The residues contributing to the H3K9me2 recognition by TD3 of hSETDB1 and their corresponding residues on DmSetDB1 are labeled with sea green. The identical residues among DmSetDB1, mSETDB1, and hSETDB1 are highlighted with black background, and the residues of the same class are shown in red. **(B)** Crystal structure of the DmSetDB1 3TD bound to the H3K9me2K14ac peptide. Residues A1-K9 of the peptide were disordered and invisible. On the left, the 3TD of DmSetDB1 is shown in the cartoon representation. On the right, electrostatic surface representation of the 3TD of DmSetDB1 bound to the H3K9me2K14ac peptide. The H3K9me2K14ac peptide is shown as a stick model and colored magenta. **(C)** 2mFo-DFc map for the H3K9me2K14ac peptide, which is contoured at 1σ by PyMOL. **(D)** Comparison of the H3K14ac residue recognition between the 3TDs of DmSetDB1 and hSETDB1 (PDB: 6BHI). **(E-F)** Comparisons of the aromatic cages in TD2 and TD3 between DmSetDB1 and hSETDB1 (PDB: 6BHI), respectively. The different residue is marked using a red dashed circle. The black dashed lines represent the hydrogen bonds formed between protein residues and the peptide. **(G-H)** Detailed interactions of the 3TD of DmSetDB1 bound to the H3K9me2K14ac peptide. Hydrogen bonds formed between protein and peptide are marked as black dashed lines. **(I)** SPR analysis of wild type (WT) Tudor domains and the F578A mutant binding to the H3K14ac peptides. The mean equilibrium dissociation constants (Kd) were determined from 3 independent experiments.

To verify this potential interaction, we determined the crystal structures of the 3TD of *Drosophila* Eggless/SetDB1 in both its apo-state and in complex with an H3 peptide containing H3K9me2 and H3K14ac modifications (H3K9me2K14ac). The 3TD of *Drosophila* Eggless/SetDB1, similar to its human counterpart, consists of three classical Tudor domains (TD1, TD2, and TD3) that adopted an antiparallel β-barrel-like structure (Figure 5B). Superposition of the structures of *Drosophila* 3TD in its apo-state and in the H3K9me2K14ac peptide-bound complex showed that the two structures overlapped very well with an RMSD value of 1.52 Å over all the Cα-atoms of the 3TD domain. However, in the *Drosophila* 3TD-H3K9me2K14ac complex, only the S10-P16 fragment of the H3K9me2K14ac peptide was observed in the electron density map, while the residues A1-K9 in the peptide were disordered and invisible (Figure 5B-5C). Further structural analysis showed that the H3K14ac peptide was positioned at the interface between the TD2 and TD3 (Figure 5B-5C). Since the 3TD of human SetDB1 interacts with H3K9me2/3 and H3K14ac doubly modified peptide (Jurkowska *et al*., 2017), we compared the structures of the *Drosophila* 3TD-H3K14ac and the human 3TD-H3K9me2/3K14ac complexes. As expected, the residues involved in H3K14ac recognition exhibited a high degree of sequence and structural similarity (Figure 5A, 5D). However, several aromatic residues involved in the H3K9me2/3 recognition by the human 3TD were not conserved in *Drosophila* 3TD. In human 3TD, the TD2 recognizes the H3K9me3 mark with an aromatic cage formed by residues Y268, W275, Y277, F297, and Y301 (Figure 5E), and the TD3 forms an aromatic cage via the residues W358, W363, and F379 to accommodate the H3K9me2 modification with an additional hydrogen bond provided by residue D382 (Figure 5F). Mutation of any of these aromatic residues in human 3TD abolished or reduced the binding affinity to the H3K9me1/2/3K14ac peptide (Jurkowska *et al*., 2017). Structural comparison revealed that W275 in human TD2 was substituted with A557 in *Drosophila* TD2, and W358 and W363 in human TD3 were replaced by S640 and L645 in *Drosophila* TD3, respectively (Figure 5E-5F). The absence of these aromatic residues in *Drosophila* 3TD disrupted the formation of complete aromatic cages, likely abrogating the recognition of H3K9 methylations.

We further analyzed the binding interface between *Drosophila* 3TD and the H3K14ac peptide. In the complex structure, the K14ac mark was stabilized through hydrophobic interactions with the side chains of F578, T584, F613, I671, and Y672 (Figure 5G). The main chains of G582 and Y672 also contributed to the recognition of the K14ac mark by forming two hydrogen bonds with the acetyl group of K14ac (Figure 5G). In addition to the acetyl group, the main chain of K14ac formed a hydrogen bond with the side chain of S675 (Figure 5G). For other residues of the peptide, the residue T11 was stabilized by forming a hydrogen bond with G582 of TD2, while G12 and G13 in the peptide made hydrogen bonding interactions with the side chains of E669 and R677 of TD3, respectively. Furthermore, the main chain carbonyl oxygen of P16 formed a hydrogen bond with the side chain of E639 of TD3 (Figure 5H). To measure the binding affinity between *Drosophila* 3TD and H3K14ac peptide, we employed surface plasmon resonance (SPR) analysis. The Kd of wild type 3TD toward H3K14ac peptide was 485.9 μM. When F578, a key residue directly interacting with the H3K14 acetyl group, was mutated to alanine (F578A), the *Drosophila* 3TD could no longer bind to the H3K14ac peptide (Figure 5I).

In summation, our comprehensive structural and binding analyses shed light on the fact that the Tudor domains of *Drosophila* Eggless/SetDB1 can indeed recognize the H3K14ac histone mark independently, without the need for H3K9 methylations, although the *in vitro* binding affinity is relatively modest.

### Disruption of the interaction between Eggless/SetDB1 and H3K14ac reduces Eggless/SetDB1 nuclear foci formation in the early embryo

To investigate the *in vivo* function of the recognition of H3K14ac by Eggless/SetDB1, we injected *Drosophila* embryos with a recently developed binder competitive inhibitor (R,R)-59 of the tandem Tudor domains of SetDB1 (Guo et al., 2021), and then performed live imaging of the distributions of the endogenously tagged GFP-SetDB1 as well as the His2Av-mRFP expressed from a transgene (Figure 6A). Injection of (R,R)-59 weakened the accumulation of GFP-SetDB1 into nuclear foci and caused cell cycle arrest in *Drosophila* syncytial embryos (Supplementary Movies S8-S9), although we could not rule out the possibility of off-target effects of the injected small molecular inhibitor.

**Figure 6.**
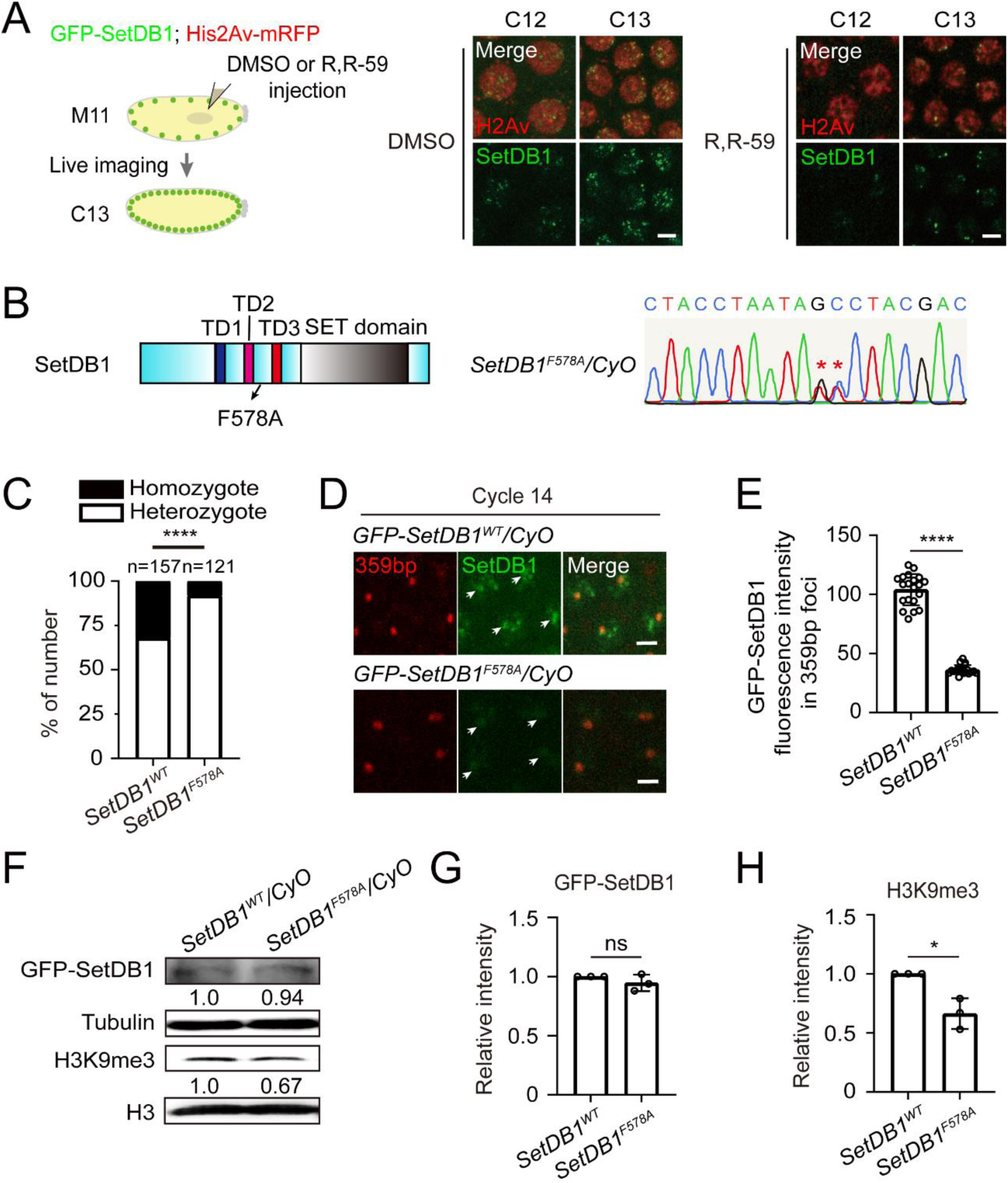
Recognition of H3K14ac by the TDs of SetDB1 is required for constitutive heterochromatin formation. **(A)** Video frames from live imaging experiments on embryos injected with DMSO or R,R-59, a small molecule targeting the TDs of SetDB1. The experiment scheme is shown on the left. The endogenously GFP-tagged SetDB1 is shown in green, and His2Av-mRFP expressed from a transgene in red. Bars: 5 μm. **(B)** Schematic of *Drosophila* SetDB1 and the key residue that is mutated in this study (left), and the Sanger sequencing validation (right). **(C)** The ratio of heterozygote/homozygote progeny from the F578A knock-in flies is greater than the expected 2:1 seen in the control. Chi-square test, ****: *p* < 0.0001. **(D)** Video frames from live imaging analysis of embryos from *GFP-SetDB1^WT^/CyO* or *GFP-SetDB1^F578A^/CyO* flies. GFP-SetDB1 is shown in green, and the 359-bp loci labeled with TALE-light is shown in red. White arrows indicate the 359-bp repeats regions. Bars: 5 μm. **(E)** Quantification of the mean GFP fluorescent intensity within the 359-bp repeats loci in cycle 14 embryos. Unpaired *t*-test, ****: *p* < 0.0001. Error bars represent the SD. **(F)** Western blot of 0-2 h embryos from *GFP-SetDB1^WT^/CyO* and *GFP-SetDB1^F578A^/CyO* flies with the indicated antibodies. **(G)** Quantification of GFP-SetDB1 western blot signals in 0-2 h embryos from *GFP-SetDB1^WT^/CyO* and *GFP-SetDB1^F578A^/CyO* flies. Unpaired *t*-test, ns: no significance. Error bar represents the SD. **(H)** Quantification of the H3K9me3 western blot signals. Unpaired *t*-test, *: *p* < 0.05. Error bar represents the SD.

To precisely disrupt the interaction between Eggless/SetDB1 and H3K14ac, we generated knock-in mutations via CRISPR-Cas9 to recode F578 to alanine (F578A) in both wild type flies and that carry the endogenously GFP-tagged *Eggless/SetDB1* alleles (Figure 6B). The homozygous F578A mutants were infertile and semi-lethal, as the ratio of heterozygotes to homozygotes in the progeny of heterozygous parents significantly deviated from the expected 2:1 seen in the wild type (Figure 6C). Even the heterozygous F578A mutants showed increased developmental defects and decreased viability. To investigate the effect of F578A mutation on the distribution of GFP-SetDB1, we collected embryos from heterozygous parents and performed live imaging analysis (Figure 6D). Compared to wild type GFP-SetDB1 that was recruited to several nuclear foci including the 359-bp repeats region in cycle 14 embryos, GFP-SetDB1^F578A^ appeared to be more diffusive, with decreased fluorescent intensity in the nuclei as well as within the 359-bp repeat loci (Figure 6D-6E, Supplementary Movies S10-S11). Western blot analysis revealed that the protein level of GFP-SetDB1^F578A^ remained comparable to that of wild type (Figure 6F-6G), suggesting that the mutation did not affect protein stability. Of note, we observed a slight decrease of H3K9me3 in embryos collected from GFP-SetDB1^F578A^ heterozygous parents (Figure 6H). These changes were reminiscent of that observed in the embryos with Gcn5 knockdown (Figure 2I-2J), indicating that disruption of Eggless/SetDB1 localization compromises the deposition of H3K9me3.

### Heterozygous *Eggless/SetDB1^F578A^* knock-in mutation compromises heterochromatin integrity

We mapped the genomic distribution of GFP-SetDB1^F578A^ in comparison to GFP-SetDB1^WT^ as well as the corresponding H3K9me3 histone modification in stage 5 embryos collected from heterozygous parental flies using ChIP-seq. The overall signals of GFP-SetDB1 and H3K9me3 after normalization were decreased in embryos carrying the GFP-SetDB1^F578A^ mutation (Figure 7A). The GFP-SetDB1 and H3K9me3 signals at the 1067 transposons and satellite elements loci that were responsive to the knockdown of Gcn5 were also markedly declined in the GFP-SetDB1^F578A^ mutant embryos (Figure 7B). At the *gypsy* and *mdg1* loci, the amounts of GFP-SetDB1 and H3K9me3 signals were significantly lower in GFP-SetDB1^F578A^ embryos compared to wild type (Figure 7C). We further performed RNA-seq to analyze the aberrant transcription of repetitive DNA elements (Supplementary Table S2). 74 out of the 122 classes of repetitive elements that were identified in the RNA-seq experiments with Gcn5 knockdown embryos manifested consistent transcriptional derepression in the GFP-SetDB1^F578A^ embryos (Figure 3A, 7D, Supplementary Table S3). We selected several transposons and validated their expression by RT-qPCR (Figure 7E).

**Figure 7.**
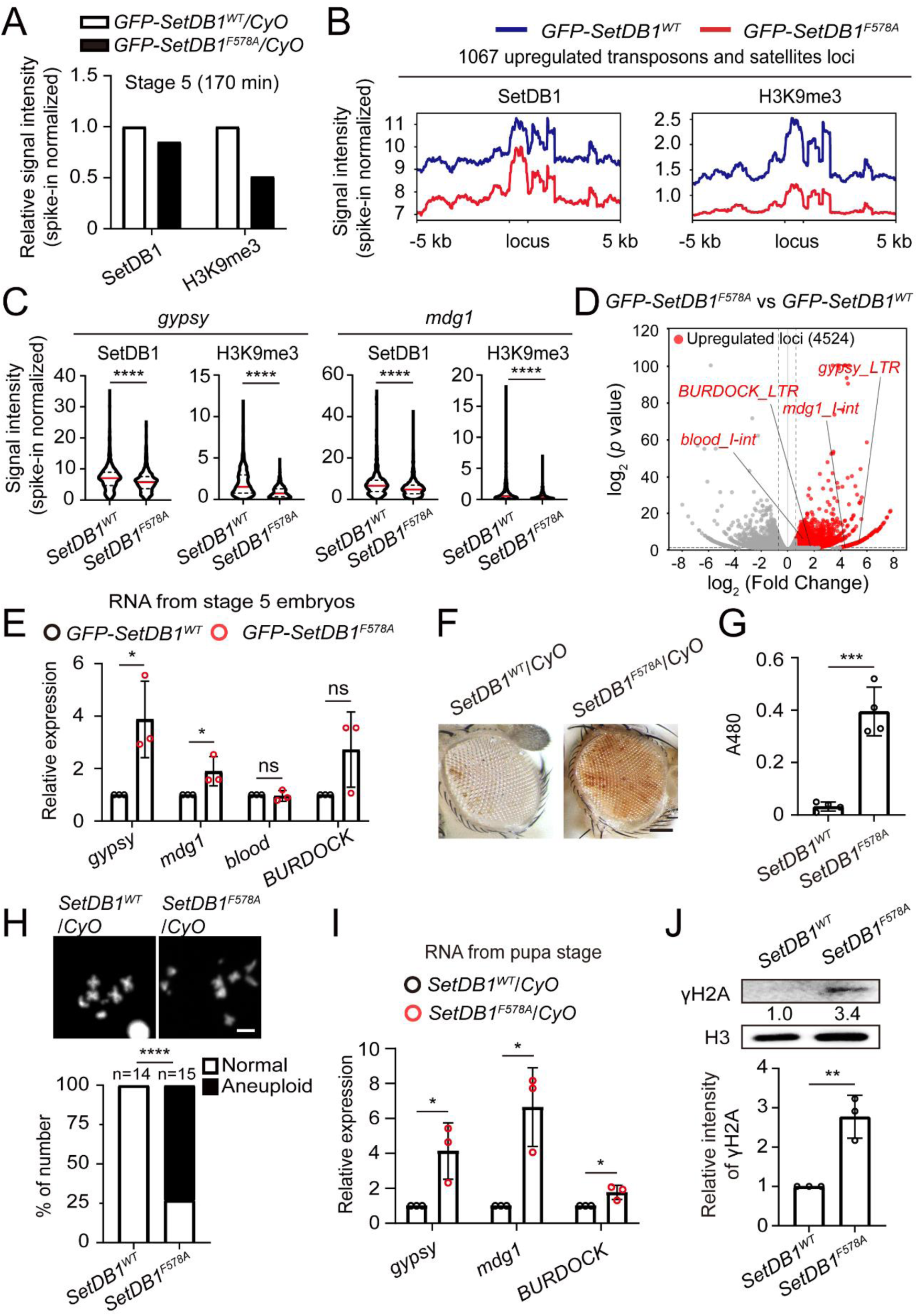
F578A knock-in of SetDB1 compromises transposon silencing and heterochromatin integrity. **(A)** Spike-in normalized ChIP-seq signals in embryos collected from *GFP-SetDB1^WT^/CyO* or *GFP-SetDB1^F578A^/CyO* flies. **(B)** Averaged ChIP-seq signals of GFP-SetDB1 and H3K9me3 around the aforementioned 1067 upregulated transposons and satellites loci in embryos collected from *GFP-SetDB1^WT^/CyO* or *GFP-SetDB1^F578A^/CyO* flies. **(C)** Violin plots of the ChIP-seq signals of GFP-SetDB1 or H3K9me3 at *gypsy* and *mdg1* genomic loci. Unpaired *t*-test, ****: *p* < 0.0001. **(D)** Volcano plot of repeats expressions in embryos (stage 5, 140 min) collected from *GFP-SetDB1^WT^/CyO* or *GFP-SetDB1^F578A^/CyO* flies. Upregulated repeat elements in the *SetDB1^F578A^* experimental group (Fold Change ≥ 1.5, *p* < 0.05) are plotted in red. **(E)** RT-qPCR analysis of the 4 *Gcn5* KD-sensitive transposons (*gypsy*, *mdg1*, *blood*, and *BURDOCK*) in embryos collected from *GFP-SetDB1^WT^/CyO* or *GFP-SetDB1^F578A^/CyO* flies. Unpaired *t*-test, ns: no significance, *: *p* < 0.05. Error bars represent the SD. **(F)** Eye pictures from the female *w^m4h^* PEV reporter flies with the indicated genotypes. Bar: 50 μm. **(G)** Pigment assay of the control flies or that carrying the *SetDB1^F578A^* knock-in mutation. Unpaired *t*-test, ***: *p* < 0.001. Error bars represent the SD. **(H)** Chromosome spreads of 3rd instar neuroblasts from control or the *SetDB1^F578A^* knock-in mutant. Chi-square test, ****: *p* < 0.0001. **(I)** RT-qPCR analysis of the indicated transposons at pupa stage. Unpaired *t*-test, *: *p* < 0.05. Error bars represent the SD. **(J)** Western blot analysis of pupae from the indicated flies writh H3 and γH2A antibodies. The quantification shows increased γH2A signal in the *SetDB1^F578A^* knock-in mutant. Unpaired *t*-test, **: *p* < 0.01. Error bar represents the SD.

The reduced deposition of H3K9me3 and transcriptional derepression of transposons suggested that the integrity of constitutive heterochromatin was compromised in the offspring of SetDB1^F578A^ heterozygous parents. Consistently, we crossed the SetDB1^F578A^ allele to the PEV reporter and observed that even heterozygous mutation of SetDB1^F578A^ resulted in increased eye pigmentation (Figure 7F-7G). Defects in constitutive heterochromatin often accompany genomic instability. In the neuroblasts of third instar larvae, the aneuploidy rate was significantly higher in SetDB1^F578A^ heterozygous mutants compared to wild type (Figure 7H). At pupa stage, many transposons such as *gypsy*, *mdg1*, and *BURDOCK* showed increased activity in SetDB1^F578A^ heterozygous mutants, and the DNA damage-associated γH2A abundance was upregulated concomitantly (Figure 7I-7J).

These results indicate that, similar to Gcn5 knockdown which erases the H3K14ac histone mark, disrupting the interaction of Eggless/SetDB1 with the H3K14ac via substitution of F578 with alanine also causes defects in the genomic distributions of Eggless/SetDB1 and H3K9me3, compromising constitutive heterochromatin integrity and organismal viability.

## DISCUSSION

### A dynamic bivalent state consisting of H3K14ac and H3K9me3 on repetitive DNA elements in *Drosophila* early embryonic development

Bivalent chromatin is tightly linked to epigenetic reprogramming and cell fate determination during animal development. In this study, by elaborating the distribution of H3K14ac and H3K9me2/3 during *Drosophila* early embryogenesis, we report a novel bivalent combination that accompanies the emergence of constitutive heterochromatin on a group of repetitive elements in the zygotic genome. H3K14ac is typically associated with active transcription and can be deposited by two acetyltransferases, HBO1 and Gcn5. *Drosophila* Chameau/HBO1 is responsible for catalyzing H3K14ac on a set of tissue-specific genes during organogenetic stage. These genes are devoid of canonical modifications such as H3K9ac, H3K27ac, and H3K4me3, and remained transcriptionally active solely through H3K14ac, as removal of H3K14ac causes decreased chromatin accessibility and RNA polymerase II occupancy on these genes (Regadas *et al*., 2021). Gcn5 is the primary acetyltransferase responsible for H3K14ac in *Drosophila.* In addition to H3K14ac, it can also catalyze H3K9ac. In Gcn5 mutant polytene chromosomes, both H3K14ac and H3K9ac are barely detectable (Carre *et al*., 2005). Consistently, H3K14ac and H3K9ac often co-occur at actively transcribing genes and regulatory elements (Karmodiya et al., 2012; Regadas *et al*., 2021). The pairing of H3K14ac with repressive histone modifications is rare, but in mouse embryonic stem cells as well as in mouse liver, the co-occupied genomic regions of H3K14ac and H3K9me3 have been detected (Jurkowska *et al*., 2017; Price et al., 2020). However, neither the establishment of this bivalent state nor its physiological function is understood.

Our results demonstrate that in *Drosophila* H3K14ac is intergenerationally transmitted from the female germline and remains present in the zygotic genome throughout the early embryogenesis. The maintenance of H3K14ac during early embryonic development is mainly mediated by Gcn5. Yet, at least four different Gcn5 complexes exist in *Drosophila*: the Spt-Ada-Gcn5 Acetyltransferase (SAGA), Ada2a-containing (ATAC), Ada2/Gcn5/Ada3 transcription activator (ADA), and Chiffon Histone Acetyltransferase (CHAT) complexes (Torres-Zelada and Weake, 2021). While it is not clear which complex controls the H3K14ac dynamics in the early embryo, we think the CHAT complex is worthy of extra attention (Torres-Zelada et al., 2022), because *Chiffon* also encodes a subunit for the Dbf4-dependent kinase (DDK) complex, which regulates cell cycle length and late replication of repetitive sequences by counteracting Rif1 activity (Seller and O’Farrell, 2018). Around the time window of the MBT (late stage 4 and stage 5) when cell cycles slow down and constitutive heterochromatin makes its first appearance, H3K14ac becomes increasingly concentrated at the apical poles of nuclei, forming bivalent genomic regions with H3K9me2/3. Our structural analysis shows that, unlike human SetDB1 (Jurkowska *et al*., 2017), *Drosophila* Eggless/SetDB1 can recognize the H3K14ac histone mark independent of the H3K9 modification status, indicating that H3K14ac nucleosomes can recruit Eggless/SetDB1 to further add H3K9me3 modification. Therefore, the establishment of the H3K14ac and H3K9me3 bivalent genomic regions in *Drosophila* early embryos appears to be a consequence rather than a cause of constitutive heterochromatin formation. Agreeingly, in stage 6 after the MBT, this bivalency is no longer maintained on repetitive sequences. The deacetylase responsible for removing H3K14ac remains unknown, but one potential candidate is Rpd3, which can counteract genomic silencing both in *Drosophila* and yeast (De Rubertis et al., 1996), and interact with Eggless/SetDB1 as well as nucleosome remodeler Mi-2 (Mugat *et al*., 2020).

### Function of H3K14ac in nuclear compartmentalization and constitutive heterochromatin formation

The restoration of epigenetic constraints in zygotic genome is central to early embryonic developmental programming. Constitutive heterochromatin, characterized by enrichment of H3K9me2/3 and HP1a, also needs to be re-established with precision to suppress repetitive DNA elements and maintain genomic stability. While the mechanisms that introduce the heterochromatic features to different repetitive sequences are complex (Yuan and O’Farrell, 2016), a key step in this process is the targeted recruitment of the H3K9 methyltransferase Eggless/SetDB1 (Seller *et al*., 2019). Eggless/SetDB1 is evolutionarily conserved, with a SUMO interacting motif (SIM) and triple Tudor domains (3TD) at the N-terminus, and a bifurcated SET domain harboring the H3K9 methyltransferase activity at the C-terminus (Fukuda and Shinkai, 2020; Ninova et al., 2020a). SetDB1 can be recruited to specific genomic locus via multiple mechanisms, and the most characterized pathway has been illustrated in mouse embryonic stem cells. The KRAB zinc finger family proteins (KRAB-zfps) recognize different retrotransposon elements via their zinc finger domain, meanwhile, bind to KAP1 via the KRAB domain. KAP1 contains a tandem PHD-bromodomain in its C-terminus, in which the PHD functions as an intramolecular E3 ligase to SUMOylate the lysine residues in its adjacent bromodomain. The SUMOylated lysines of KAP1 are in turn recognized by the SIM of SetDB1, mediating its selective enrichment on retrotransposons (Fukuda and Shinkai, 2020). While the similar DNA-binding protein mediated recruitment of Eggless/SetDB1 is less understood in *Drosophila*, a pioneer factor GAF that can specifically binds to GA repeats, is required for the establishment of constitutive heterochromatin on AAGAG satellite sequences (Gaskill *et al*., 2023). How GAF interacts with Eggless/SetDB1 remains unclear. Another well-studied pathway for Eggless/SetDB1 recruitment is illustrated in *Drosophila* ovaries. Relying on its interacting small RNAs (piRNAs), the Argonaute family protein Piwi is selectively engaged with nascent transcripts from the corresponding transposons. Piwi forms a complex with Asterix/Gtsf1, Panoramix/silencio, Nxf2, Nxt1, Ctp, and the SUMO E3 ligase Su(var)2-10, which induces local SUMOylation on yet-to-be-identified substrates to recruit Eggless/SetDB1 (Batki *et al*., 2019; Mugat *et al*., 2020; Ninova *et al*., 2020b; Zhao *et al*., 2019).

Functional analyses have revealed that the known Eggless/SetDB1 recruitment mechanisms only guide heterochromatin formation on a small portion of repetitive elements (Fabry *et al*., 2021; Gaskill *et al*., 2023). Our study reports a new mode of Eggless/SetDB1 recruitment, in which the active histone modification H3K14ac can directly attract Eggless/SetDB1 via its N-terminal 3TD domain. Many repetitive elements seem to be regulated by this mechanism in the early embryonic development, as disrupting the interaction between 3TD and H3K14ac causes a significant reduction of Eggless/SetDB1 binding and the H3K9me3 level on these loci. It is not immediately clear how H3K14ac-mediated Eggless/SetDB1 recruitment achieves selectivity during constitutive heterochromatin formation. Phase separation mechanisms, driven by multivalent weak interactions, might underlie the H3K14ac-Eggless/SetDB1 dependent heterochromatic compartmentalization. Eggless/SetDB1 contains many low complexity sequences and is predicted to have the capacity to undergo phase separation. In addition, Eggless/SetDB1 carries several PXVXL motifs that can interact with HP1a, which has been reported to undergo liquid-liquid phase separation that drives the formation of heterochromatic domains in the early embryos (Strom et al., 2017). Given that many repetitive elements such as the 359-bp repeats lie within the compacted chromatin before the appearance of other heterochromatic features (Shermoen et al., 2010), we reason that their DNA sequences may have certain intrinsic properties (e.g. binding linker histones or positioning nucleosomes) that allow for the assembly of nucleosomes at high density. Even though the distribution of H3K14ac is ubiquitous, the higher nucleosome density results in a higher local concentration of H3K14ac that would recruit more Eggless/SetDB1. Once the phase separation threshold is exceeded, Eggless/SetDB1 undergoes liquid-liquid demixing, introducing H3K9me3 on adjacent nucleosomes and concentrating HP1a. This process breaks symmetry and initiates nuclear compartmentalization, eventually leading to the establishment of constitutive heterochromatic regions (Figure 8).

**Figure 8.**
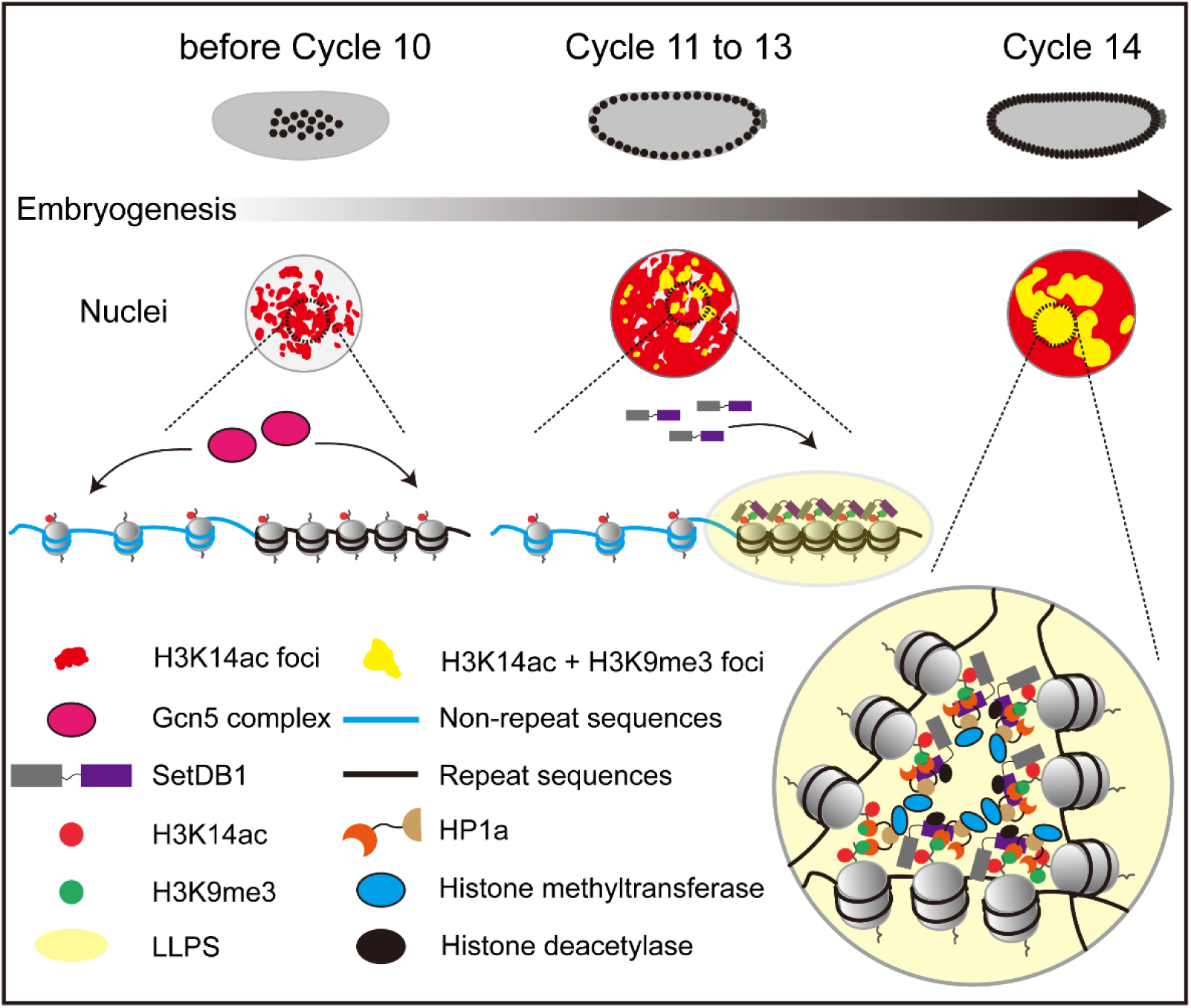
A proposed model illustrating how SetDB1 condensates on the H3K14ac-clustered repetitive sequences to participate in constitutive heterochromatin establishment.

### Limitations of the study

In this study, we showed that the maternally inherited H3K14ac recruits Eggless/SetDB1 via its tandem Tudor domain to a group of repetitive elements for the re-establishment of constitutive heterochromatin. However, the widespread distribution of H3K14ac, typically associated with active transcription, raises the question of how H3K14ac-mediated Eggless/SetDB1 recruitment selectively silences repetitive elements. Although we propose that certain repetitive elements with higher nucleosome density, such as the 359-bp repeats, could drive local phase separation of Eggless/SetDB1, this hypothesis requires experimental validation and further investigation.

## STAR★Methods

### Key resources table

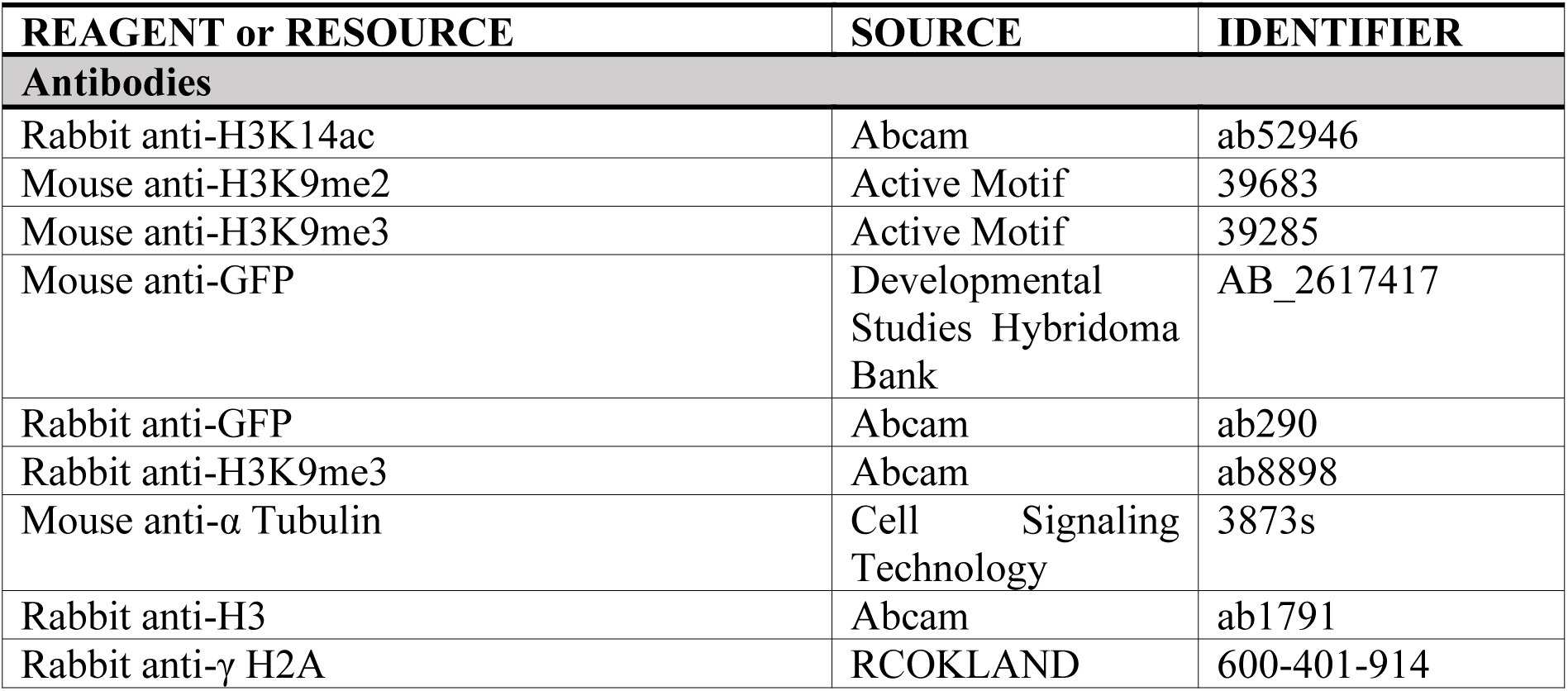
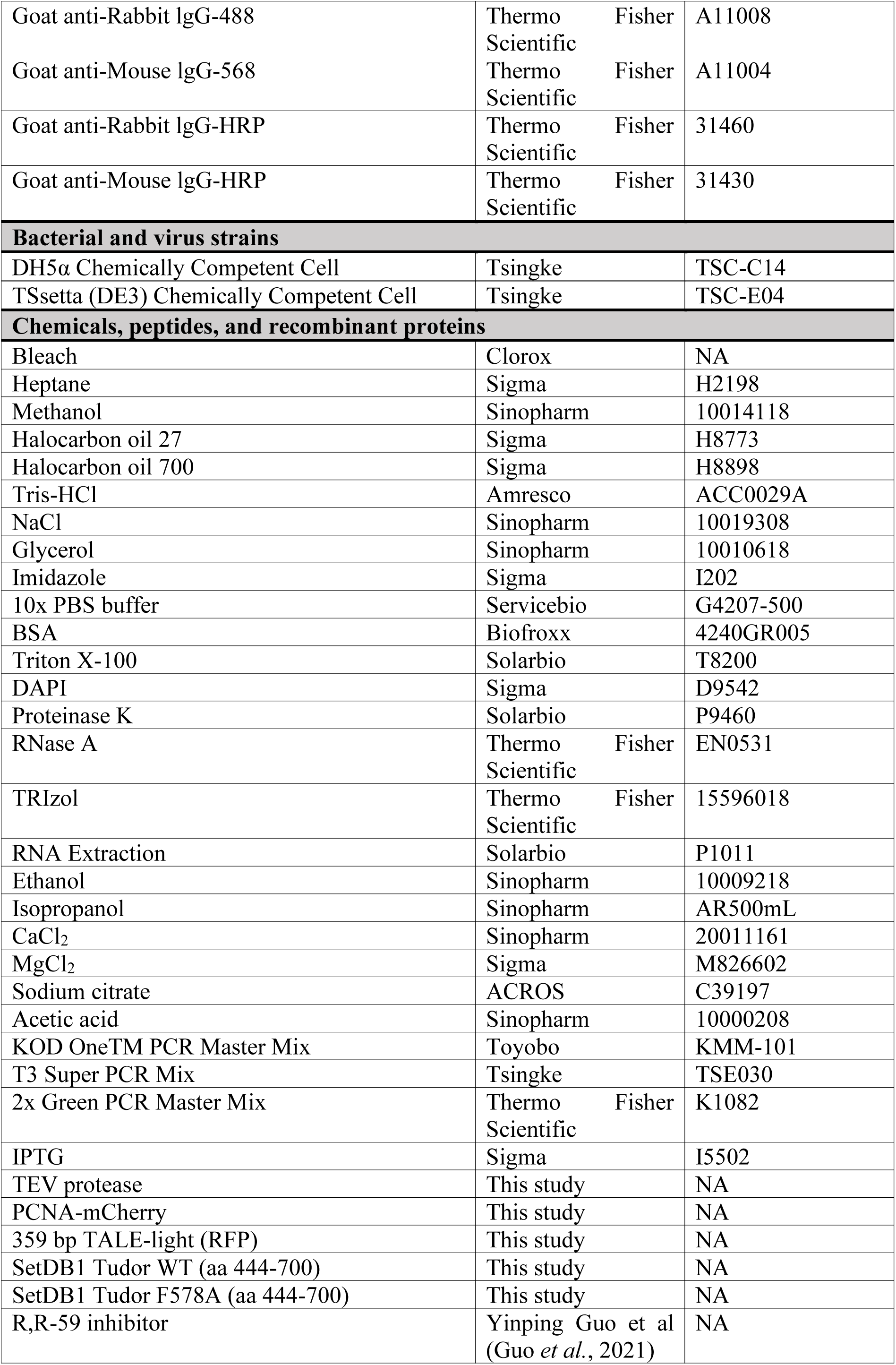
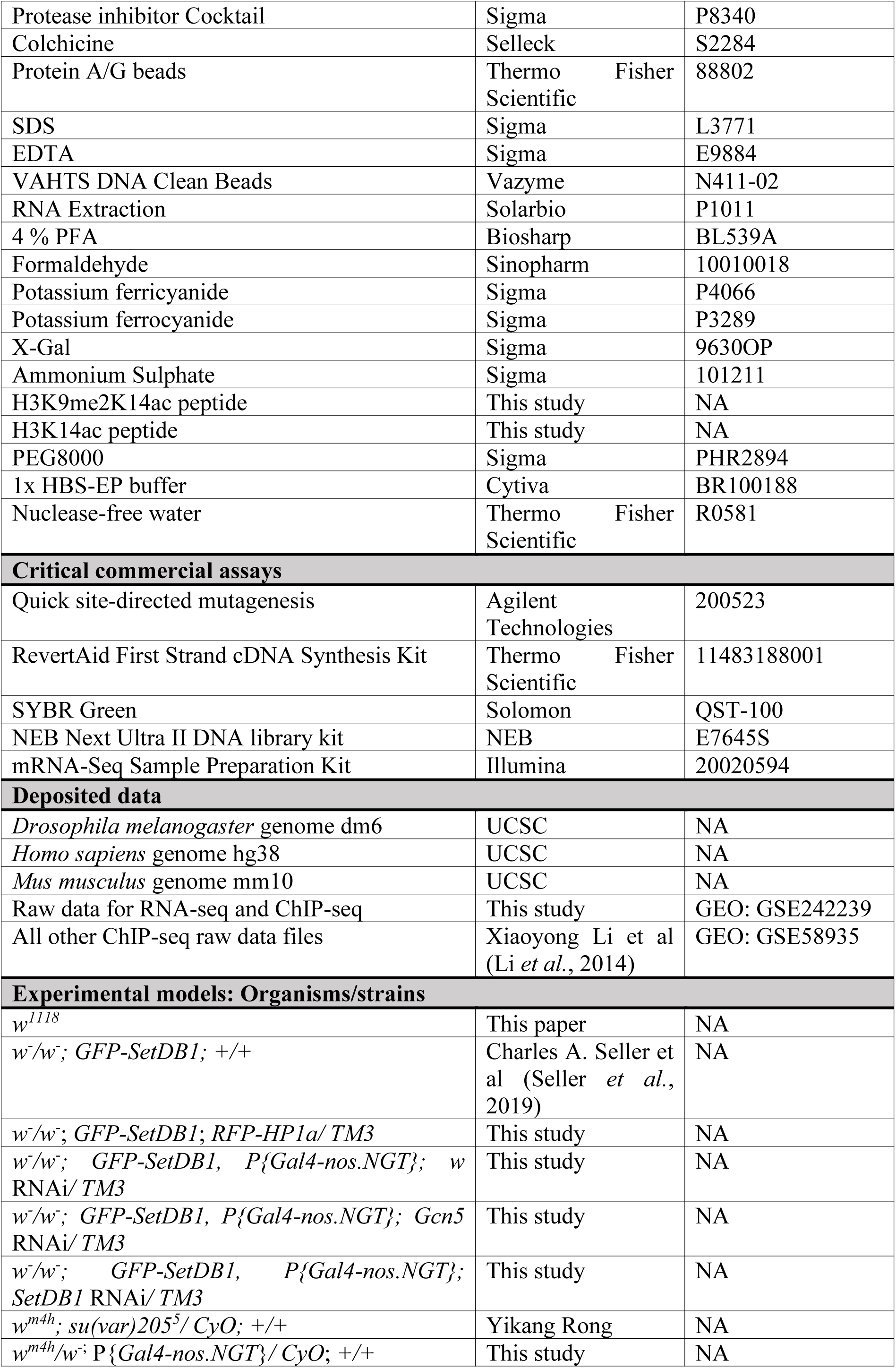
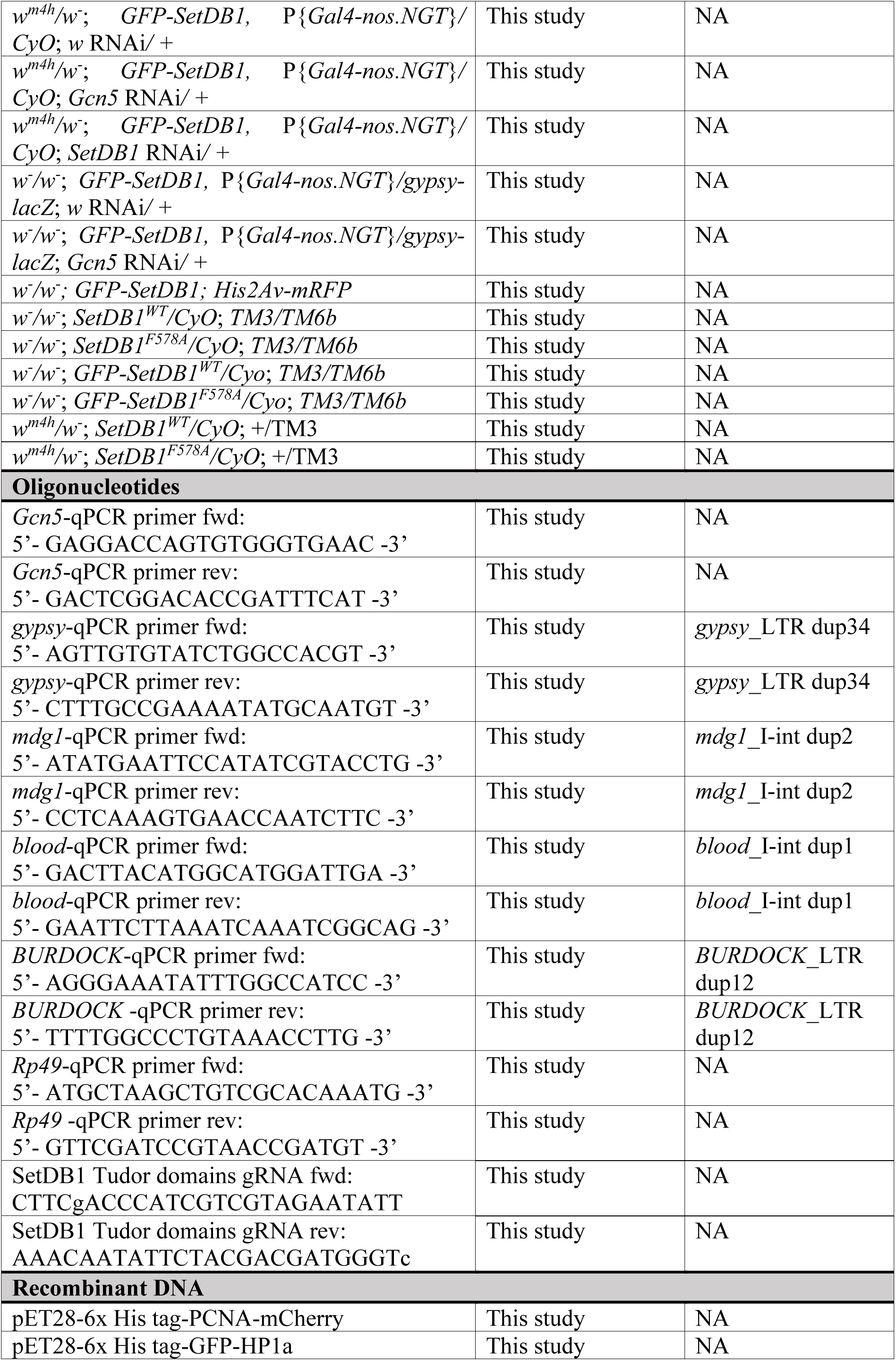
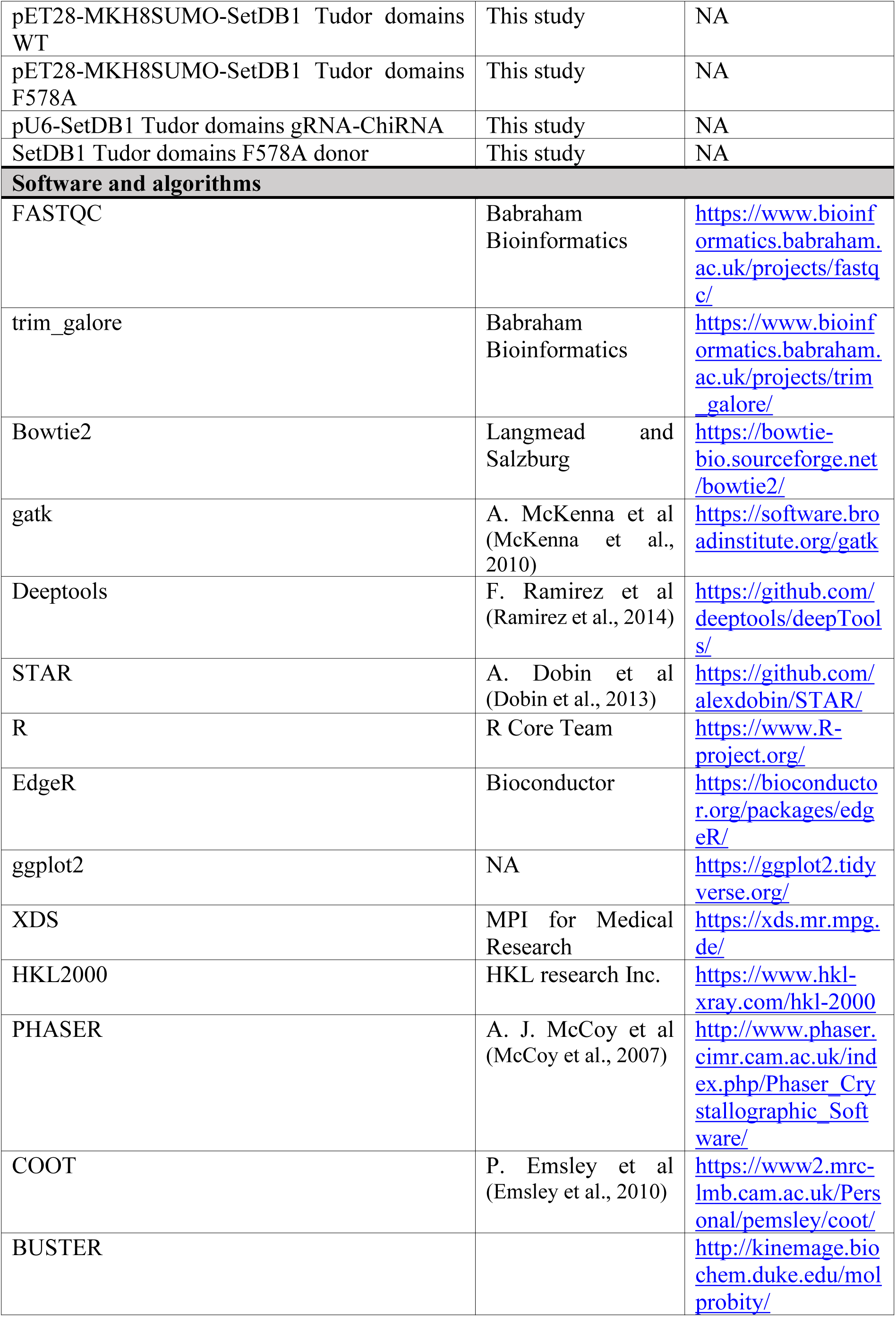
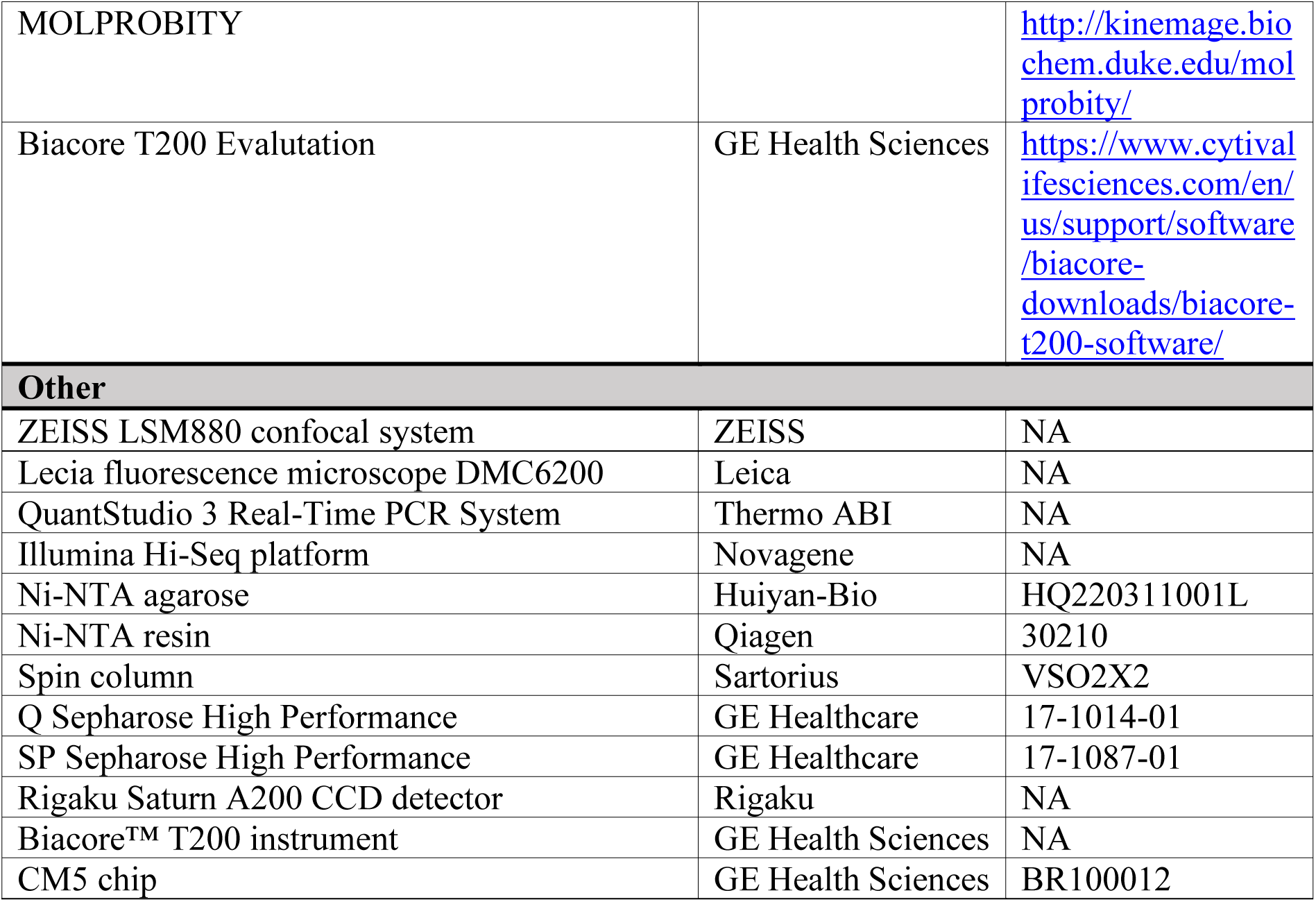

### Resource availability

#### Lead contact

Further information and requests for resources and reagents should be directed to and will be fulfilled by the lead contact, Kai Yuan.

#### Materials availability

All fly strains generated in this study are available from the lead contact without restriction.

#### Experimental model and study participant details

*Drosophila melanogaster* strains were cultured under standard laboratory conditions at 25°C. Samples were prepared as described in the Method details. All fly strains are listed in the Key resources table.

### Method details

#### Protein expression and purification

The recombinant fusion proteins, including PCNA-mCherry and 359-bp TALE-light-RFP, were purified as previously described (Seller *et al*., 2019; Yuan and O’Farrell, 2016). Briefly, the corresponding DNA sequences were cloned into pET28-His-tag vector and transformed into the TSsetta (DE3) *Escherichia coli* cells. Protein expression was induced by treating the cultures with 0.5 mM isopropyl-β-d-thiogalactopyranoside (IPTG) overnight at 20°C. The harvested cells were then resuspended in a lysis buffer containing 20 mM Tris-HCl (pH 7.5), 500 mM NaCl, and 10 mM Imidazole. After pressure crushing, the supernatant was collected by centrifugation at 12000 rpm for 30 minutes. The proteins were purified from the cleared lysate using Ni-NTA agarose (Huiyan-Bio) and concentrated using a spin column (Sartorius). The concentrations of the purified proteins were determined to be 5 mg/mL, and 2.5 mg/mL, respectively.

For the *Drosophila* SetDB1-Tudor domains (aa 444-700), we generated a sub-clone into a modified pET28-MKH8SUMO vector to produce an N-terminal His-tagged and SUMO-tagged fusion protein. The SetDB1-Tudor F578A mutant was generated using SetDB1-Tudor (aa 444-700) as a template through Quick site-directed mutagenesis (Agilent Technologies). All fusion proteins were overexpressed using (DE3) *Escherichia coli* cells by inducing with 0.25 mM IPTG at 16°C overnight. The cells were harvested and resuspended in a lysis buffer containing 20 mM Tris-HCl (pH 7.5), 500 mM NaCl, 5% glycerol, and 5 mM Imidazole. After sonication at 4°C, the supernatant was collected by centrifugation at 10000 rpm for 1 hour and further purified using Ni-NTA resin (Qiagen). The His and SUMO tags were removed using TEV protease (purified in-house) at an approximate molar ratio of 1:20 at 4°C overnight. The treated proteins were further purified using an ion exchange column (GE Healthcare). Finally, the purified proteins were stored at a concentration of 7 mg/mL by snap freezing in liquid nitrogen.

#### Micro-injection and live imaging

All parent flies were housed in a cage and the embryos were collected using a juice plate at room temperature for a duration of 30 minutes. Subsequently, the embryos were allowed to age for an additional 1.5 hours. To dechorionate the harvested embryos, they were treated with 50% fresh bleach for approximately 2.5 minutes. Retained bleach was then washed away using water, and the embryos were carefully aligned in columns on a plate. The arranged embryos were glued to a homemade coverslip and dried in a box containing CaCl_2_. Finally, a mixture of Halocarbon oil 27 and Halocarbon oil 700, in a volume ratio of 1:1, was added to the homemade coverslip.

The micro-injection was performed following a previously established protocol (Tang et al., 2020). For live labeling, recombinant PCNA-mCherry purified from *Escherichia coli* cells was used at concentration of 5 mg/mL. The SetDB1 Tudor domains inhibitor (R,R)-59, gifted from Shengyong Yang, was used at a concentration of 400 mM. The rabbit anti-H3K14ac (Abcam) and rabbit anti-H3K9me3 (Abcam) antibodies were injected at concentrations of 2.5 mg/mL and 1.5 mg/mL, respectively. Embryos were then imaged using the ZEISS LSM880 confocal system, equipped with a 63x Plan-Apochromat 1.4 NA oil objective, at room temperature. Image stacks were acquired with a 1 μm interval over a 4 μm range.

#### Immunofluorescence

The staining procedure for embryos and ovaries was described in previous studies (Tang *et al*., 2020; Yuan and O’Farrell, 2016). Briefly, embryos were collected on juice agar plates, dechorionated for approximately 2 minutes in 50% bleach, and fixed in a methanol-heptane solution (1:1) for 5 minutes. Prior to immunostaining, the embryos underwent a gradual rehydration process using a methanol concentration gradient (5 minutes each in 80%, 50%, and 20%, followed by 10 minutes in PTA). PTA is composed of 1x PBS supplemented with 0.1% Triton X-100 and 0.02% azide. Subsequently, the embryos were blocked in PBTA (PTA with 1% BSA) for 30 minutes at room temperature and incubated with primary antibodies in PBTA overnight at 4°C. The primary antibodies used included rabbit anti-H3K14ac (Abcam, 1:40), mouse anti-H3K9me2 (Active Motif, 1:200), mouse anti-H3K9me3 (Active Motif, 1:200), and mouse anti-GFP (DSHB, 1:30). Following three 5-minute washes in PBTA, the embryos were incubated with suitable fluorescently labeled secondary antibodies (Thermo Fisher Scientific, 1:400) in the dark for 1 hour.

Ovaries and testes were dissected from 7-day-old females and males, respectively, in 1x PBS. The dissected tissues were fixed in 4% PFA at room temperature for 1 hour, rinsed three times in PBST (1x PBS with 0.3% Triton X-100), and blocked in PBST with 5% BSA for 1.5 hours at room temperature. The tissues were then incubated with primary antibodies diluted in PBST with 5% BSA overnight at 4°C. The primary antibodies used in this step were rabbit anti-H3K14ac (Abcam, 1:200) and mouse anti-H3K9me3 (Active Motif, 1:200). After three 10-minute washes in PBST, the samples were incubated with appropriate fluorescently labeled secondary antibodies (Thermo Fisher Scientific, 1:400) in the dark for 1 hour. All samples were stained with DAPI and washed three times with PBST before being mounted in SlowFade Diamond mountant (Thermo Fisher Scientific). Imaging of all samples was performed using the ZEISS LSM880 confocal system.

#### Western blot

Embryos were collected from the indicated genotype parents for a duration of 120 minutes. The pupae at stages P10 to P14 were collected for analysis in this study. To prepare the samples, all specimens were first frozen using liquid nitrogen and then grounded in a 2% SDS sample buffer containing protease inhibitors (Sigma) on ice. Subsequently, the samples were sonicated on ice and boiled for 10 minutes. For western blot, the following antibodies were used: mouse anti-tubulin (CST, 1:2000), rabbit anti-GFP (Abcam, 1:1000), rabbit anti-H3 (Abcam, 1:2000), rabbit anti-H3K14ac (Abcam, 1:1000), rabbit anti-H3K9me3 (Abcam, 1:1000), and rabbit anti- γH2A (ROCKLAND, 1:2000).

#### RT-qPCR

Total RNA was extracted from collected stage 5 embryos (approximately 140 minutes of aging) using TRIzol reagent (Thermo Fisher Scientific). The extracted RNA was then reverse transcribed into complementary DNA (cDNA) using the RevertAid First Strand cDNA Synthesis Kit (Thermo Fisher Scientific). Quantitative PCR (qPCR) was performed with SYBR Green master mix (Solomon) on the QuantStudio 3 Real-Time PCR System (Thermo Fisher Scientific). The primers for target transposons were designed based on the upregulated transposons’ sequences in our RNA-seq data. To ensure accurate quantification, the expression levels of each target gene or repeat were normalized to the reference gene *Rp49*. The qPCR primers utilized in this study are provided in the key resources table.

#### Chromosome spreading

3rd instar larvae at the wandering stage were gently picked from the feeding bottle and transferred to a saline solution containing 0.7% NaCl. Subsequently, the larval brains were carefully dissected and rinsed with saline at room temperature for a duration of 1 to 5 minutes. Afterward, the dissected brains were transferred to 2 mL of saline supplemented with 100 μL of a 10 mM colchicine solution (Selleck). The brains were then incubated at a temperature of 25 °C for 1.5 hours. Following the incubation, the brains were transferred to a hypotonic solution consisting of 50 μL of 0.5% sodium citrate and incubated at room temperature for 8 minutes. Subsequently, the brains were moved to a freshly-prepared 50 μL fixative solution composed of acetic acid, methanol, and water in a ratio of 11:11:2. The fixation process continued until the brains became transparent, which typically took approximately 20 seconds. Once fixed, the brains were gently transferred to a drop of 45% acetic acid placed on a coverslip. An inverted microscope slide was carefully lowered onto the coverslip to pick it up, and slight pressure was applied to the coverslip to remove excess liquid. The slide was rapidly frozen using liquid nitrogen, causing the coverslip to detach with the aid of a surgical blade. Subsequently, the slide containing the brains was dehydrated in ethanol for at least 15 minutes at a temperature of −20 °C, and then allowed to air dry on paper towels. Finally, the remaining brain tissues on the slide were stained with DAPI before being imaged using confocal microscopy.

#### ChIP-seq and analysis

All embryo samples were collected under standard conditions at 25 °C. Following a 20 minutes collection from population cages, the embryos were aged for 90, 120, and 150 minutes to target stages 4, 5, and 6 respectively. The embryos were dechorionated by treatment with 50% bleach. Subsequently, the embryos were preserved in a 1x embryo buffer consisting of 0.7% NaCl and 0.05% Triton X-100. Hand sorting was performed in a 6 cm dish under a light microscope to exclude embryos younger or older than the target stages. The sorted embryos were snap frozen in liquid nitrogen and stored at −80 °C. Chromatin immunoprecipitation (ChIP) was performed following a previously reported protocol (Li *et al*., 2014). Approximately 5000 *w* knockdown or *Gcn5* knockdown stage 5 embryos, and 3000 stage 5 embryos from *GFP-SetDB1^WT^/CyO* and *GFP-SetDB1^F578A^/CyO* parents were utilized in a single ChIP-seq experiment. To investigate the dynamic distributions of H3K14ac, H3K9me2, and H3K9me3 throughout early embryo development, a total of 12000 *w^1118^* embryos at stage 4 (110 minutes), 6000 embryos at stage 5 (140 minutes), and 3000 embryos at stage 6 (170 minutes) were utilized in the ChIP-seq experiments.

All frozen embryos were grounded in a 1% formaldehyde solution and crosslinked for approximately 7 minutes at room temperature with shaking (the total processing time did not exceed 10 minutes). The fixation was quenched by adding a final concentration of 350 mM glycine and incubating for 10 minutes at room temperature with shaking. The nuclei were then pelleted for 2 minutes at 4 °C, 2000 rpm, and washed twice with ice-cold buffer A1 (60 mM KCl, 15 mM NaCl, 4 mM MgCl_2_, 15 mM HEPES, pH 7.6, 0.5% Triton X-100, 0.5 mM DTT, and 10 mM sodium butyrate). Next, the nuclei pellets were resuspended in 1 mL of ice-cold lysis buffer 1 (50 mM HEPES, 140 mM NaCl, 1mM ethylenediaminetetraacetic acid, 1% Triton X-100, 0.1% Na-Deoxycholate, 1 mM phenylmethylsulfonyl fluoride, 1:1000 cocktail protease inhibitors). The nuclei were then transferred to lysis buffer 2 (0.1% SDS and 0.5% N-lauroylsarcosine in fresh lysis buffer 1) at a density of 1000 embryos per 250 μL. The nuclei were then incubated in lysis buffer 2 at 4 °C for 2 hours with gentle rotation at 60 rpm. The chromatin obtained was fragmented into suitable sizes ranging from 200 to 300 bp using a Qsonica instrument (Duty Cycle - 10%; Intensity - 5; Cycles per Burst - 200; Time - 4 minutes). Prior to performing chromatin immunoprecipitation, we mixed the chromatin (5 μg) from each sample with a comparable amount of spike-in chromatin (1 μg) isolated from 293T or CG1 cells. We used 5 μg of antibodies in each chromatin immunoprecipitation reaction, including anti-H3K14ac (Abcam), anti-H3K9me2 (Abcam), anti-H3K9me3 (Abcam), and anti-GFP (Abcam). The antibody-chromatin complexes were captured by incubating with Protein A/G beads (Thermo Fisher Scientific). Afterward, the complexes underwent three washes and were eluted with a buffer containing 1% SDS, 10 mM EDTA, and 10 mM Tris-HCl (pH 8.0) at 65 °C for 30 minutes. To remove any remaining RNA, RNase A (Sigma) was applied to each sample, and the DNA fragments were released from the antibody-chromatin complexes using Proteinase K (Sigma). The samples were then cleaned up through VAHTS DNA Clean Beads (Vazyme) extraction and eluted in TE buffer. Sequencing libraries were prepared using the NEB Next Ultra II DNA library kit and sequenced on the Illumina Hi-Seq platform (Novagene).

For data processing, we checked the quality of the raw data using FASTQC (version 0.11.9). Adaptor sequences and low-quality reads were removed using trim_galore (version 0.6.0). Sequenced reads from embryos were mapped to the *Drosophila melanogaster* genome dm6, while spike-in reads were mapped to the *Homo sapiens* genome hg38 or *Mus musculus* genome mm10 using Bowtie2 (version 2.3.5.1) with default parameters. Duplicate reads were removed using MarkDuplicates from the gatk package (version 4.1.4.1). Normalization of the data using spike-in was performed as previously described (Niu et al., 2018; Orlando et al., 2014). First, we combined the number of mapped reads from both the *Drosophila* and spike-in species genomes to obtain the total mapped reads for each sample (Supplementary Table S4). We denoted the spike-in scale factor as α, the abundance of histone modification or protein as β, the percentage of other species reads in total mapped reads in input as γ, the number of spike-in species mapped reads (in millions) in the immunoprecipitant as Ns, and the number of *Drosophila* mapped reads (in millions) in the immunoprecipitant as Nd. We calculated the spike-in scale factor as α = γ/Ns and the modification or protein binding level as β = Nd * α. Relative signal levels were determined as the ratio of the experimental group to the control group. Next, we multiplied the number of mapped *Drosophila* reads by the scale factor to normalize the total read count in different samples using deeptools (version 3.5.0) function bamCoverage with the command-line options ‘--scaleFactor α -bs 200’. Heatmaps were generated using the deeptools function plotHeatmap with z-score normalized BigWig files. Correlation maps were generated using the deeptools function plotCorrelation with the command options ‘-c spearman -p heatmap --colorMap bwr’.

#### RNA-seq and analysis

The RNA extraction from stage 5 embryos was performed using TRIzol following the manufacturer’s protocol. Approximately 200-300 embryos were collected at room temperature and incubated in 500 μL of TRIzol. The embryos were gently grounded at room temperature for 10 minutes, followed by the addition of 100 μL of RNA Extraction (Solarbio) and vortexed for 15 seconds. The samples were then incubated at room temperature for an additional 10 minutes. Following the incubation, the samples were centrifuged at 10000 g at 4 °C for 15 minutes, and the supernatant was collected. Isopropanol was added in equal volume to precipitate the RNA, which was subsequently washed three times with 75% ethanol. Finally, the RNA was resuspended in nuclease-free water (Thermo Fisher Scientific). For library preparation and sequencing, the extracted total RNA was processed using the mRNA-Seq Sample Preparation Kit (Illumina). The sequencing was conducted on an Illumina Hiseq platform (Novagene) adhering to the standard protocols.

The resulting RNA-seq data was processed as follow. Initial read cleaning was performed using trim_galore (version 0.6.0) with default parameters. Following cleaning, the reads from each RNA-seq sample were mapped to the dm6 genome assembly obtained from UCSC, using STAR (version 2.5.3a). The alignment parameters used for this process were set as follow: ‘-- outFilterMismatchNoverLmax 0.04 --outSAMtype BAM SortedByCoordinate -- outFilterMultimapNmax 500 --outMultimapperOrder Random --outSAMmultNmax 1’. Repeats expression was quantified using featureCounts (version 1.6.5) based on the annotation files from UCSC. To label different repeats loci, we assigned unique identifiers to the repeats loci in the last column of the dm6_repeat.gtf file, such as ‘repeat_locus_1’. The unique repeats were identified using featureCounts (version 1.6.5) with the parameter ‘-a dm6_repeat.gtf -o clous.counts -M -g repeat_locus’. The identification of 1067 transposon and satellite repeats loci was accomplished using the R package dplyr (version 4.1.3). Differential expression analysis was conducted using the R package EdgeR (version 3.14.0). The differentially expressed repeats were visualized using the R package ggplot2 (version 0.9.3) to generate volcanoplots.

#### Position-effect variegation assay

The PEV assay was conducted following a previously reported procedure (Elgin and Reuter, 2013; Gu and Elgin, 2013). Briefly, female flies of the indicated genotypes were crossed with PEV reporter males, including *nos-Gal4*, *GFP-SetDB1, nos*-*Gal4*; UAS-*w* RNAi, *GFP-SetDB1, nos*-*Gal4*; UAS-*Gcn5* RNAi, *GFP-SetDB1, nos*-*Gal4*; UAS-*SetDB1* RNAi, *SetDB1^WT^/CyO* and *SetDB1^F578A^/CyO*. The resulting female offspring was collected and subjected to analysis. Adult eye imaging was performed using a Lecia fluorescence microscope DMC6200. For pigment quantification, the heads of 10 female adult files were dissected and rinsed in 1x PBS. Subsequently, the heads were homogenized in 0.1% HCl in methanol using a grinding rod. The homogenate was centrifuged at 12000 rpm for 10 minutes and the resulting supernatant was collected. The emission spectra of the supernatant were evaluated at 480 nm using a spectrophotometer.

#### X-Gal staining

The staining protocol was described previously (Batki *et al*., 2019; Mugat *et al*., 2020). In short, the ovaries were dissected from 4-5 days old female flies in cold 1x PBS. The dissected ovaries were then fixed in 2% formaldehyde/PBS at room temperature for 10 minutes, followed by three washes with 1x PBS. Subsequently, the fixed samples were incubated with a staining solution containing 1x PBS (pH 7.5), 1 mM MgCl_2_, 150 mM NaCl, 3 mM potassium ferricyanide, 3 mM potassium ferrocyanide, 0.1% Triton X-100, and 0.1% X-Gal at room temperature overnight.

#### Protein crystallization

The purified SetDB1 3TD proteins (7 mg/mL) were subject to crystallization using the sitting drop vapor diffusion method at a temperature of 18 °C. This was achieved by combining 0.5 μL of the complex sample with an equal volume of the reservoir solution. The apo crystals were successfully obtained under the following condition containing 1.6 M Ammonium Sulphate, 0.01 M MgCl_2_, and 0.1 M Tris at a pH of 8.5. To obtain the SetDB1 3TD-peptide complex crystal, the purified protein was mixed with different H3K9me1/2/3K14ac or H3K14ac peptides at a molar ratio of 1:3. The mixture was then incubated at room temperature for 20 minutes, followed by crystallization using the aforementioned sitting drop vapor diffusion method. The DmSetDB1-H3K9me2K14ac complex crystals were obtained in a buffer comprising 15% PEG8000 (w/v), 0.2 M MgCl_2_, and 0.1 M Tris (pH 8.5). Prior to flash-freezing, the crystals were safeguarded by immersing them in a cryoprotectant composed of the reservoir solution with an additional 20% glycerol.

#### Data collection and structure determination

X-ray diffraction data were collected at 100K using the Rigaku Saturn A200 CCD detector. The collected data were subsequently processed utilizing XDS (Kabsch, 2010) and HKL2000 software (Otwinowski and Minor, 1997). Structures were solved by molecular replacement using PHASER (McCoy *et al*., 2007) with PDB entry 6BHD and 6BHH as search templates. Manual model rebuilding was performed using COOT software (Emsley *et al*., 2010), and structural refinement was executed using BUSTER (version 2.10.0) (Smart et al., 2012). The resulting structures were validated using MOLPROBITY (Davis et al., 2004). Detailed statistics regarding the diffraction data and structure refinement can be found in Supplementary Table S5.

#### Surface plasmon resonance assays

SPR assays were conducted using the Biacore™ T200 instrument (GE Health Sciences Inc.). Firstly, the CM5 chip was activated following the protocol provided by GE Health Sciences Inc. The SetDB1-Tudor domain protein was then immobilized onto the CM5 chip at immobilization levels of approximately 4000 RU. For binding analysis, a five-point concentration series of the peptide was prepared through serial dilutions ranging from 6.175 μM to 500 μM in a 1x HBS-EP buffer (0.01 M HEPES, pH 7.4, 0.15 M NaCl, 3 mM EDTA, 0.005% P20). Each peptide sample was flowed at a rate of 75 μL min^−1^, and single-cycle kinetic analysis was conducted with an on-time of 60 s and an off-time of 120 s. Curve fitting and Kd determination were performed using the Biacore T200 Evaluation software (GE Health Sciences Inc.). The results were obtained from three independent replicates.

#### Quantification and statistical analysis

For quantification of fluorescence signals, images were analyzed and processed using ImageJ (https://imagej.nih.gov/ij/) and ZEN blue (https://www.zeiss.com/microscopy/en/products/software/zeiss-zen.html) software. All statistical analyses were completed in Graphpad Prism 9 (https://www.graphpad.com/scientific-software/prism/) software. Detailed descriptions of statistical analyses can be found in the figure legends. All data are presented as mean ±SD.

#### Additional resources

This study did not generate any additional resources.

#### Data and Code Availability

All sequencing data were deposited at NCBI Gene Expression Omnibus (GEO) and are publicly available as of the date of publication. Accession numbers are listed in the key resources table. The crystal structures of the Eggless/SetDB1 3TD in apo form and in complex with H3K9me2K14ac peptide were deposited in the Protein Data Bank with accession codes 7UW8 and 7UVE, respectively.

Any additional information required to reanalyze the data reported in this paper is available from the lead contact upon request.

## ACKNOWLEDGEMENTS

We gratefully acknowledge Drs. Patrick H. O’Farrell, Shengyong Yang, Yikang Rong, Zhouhua Li, Yang Yu, Jianye Zang, Su Qin, Guanjun Gao, the Developmental Studies Hybridoma Bank, the Bloomington *Drosophila* Stock Center, and TsingHua Fly Center for inspiring discussions or reagents. We would also like to acknowledge the assistance of Huan Liu in the crystallization and Fengling Li in the SPR analysis. This project has been supported by the National Natural Science Foundation of China (grants 32170821, 92153301, and 32370821 to K.Y, 32101034 to F.C, 31770834 to K.L), National Key Research and Development Program of China (2021YFC2701200), Department of Science & Technology of Hunan Province (grants 2023RC1028, 2021JJ10054, and 2023SK2091 to K.Y).

## Author contributions

Conceptualization: K.L. and K.Y.; Methodology: R.T., M.Z., Y.C., Z.J., X.F., A.D., S.M., F.C., K.L., K.Y.; Validation: Y.C., X.F., J.Z., L.L., S.M.; Software: R.T., S.M.; Formal Analysis: R.T., M.Z., K.Y.; Investigation: R.T., M.Z., Y.C., Z.J., K.Y.; Resources: J.M., K.L., K.Y.; Data Curation: R.T., M.Z., Z.J.; Writing-Original Draft: R.T.; Writing-Review & Editing: Y.C., J.M., K.L., K.Y.; Visualization: R.T., K.L., K.Y.; Supervision: K.L., K.Y.; Project Administration: K.Y., L.L.; Funding Acquisition: K.L., K.Y.

## Supplemental Tables

Table S1. The differentially expressed repeats in *Gcn5* KD embryos versus the *w* KD control, related to Figure 3 and 4.

Table S2. The differentially expressed repeats in embryos carrying GFP-SetDB1^F578A^ versus that carrying GFP-SetDB1^WT^, related to Figure 7.

Table S3. The upregulated 74 classes of transposons & satellites.

Table S4. The details of the spike-in analysis of ChIP-seq data in this study.

Table S5. Statistics of the diffraction data and the structure refinement.

## Supplemental Movies

Movie S1. Control embryo microinjected with PBS.

Movie S2. The embryo microinjected with H3K9me3 antibody.

Movie S3. The embryo microinjected with H3K14ac antibody.

Movie S4. GFP-SetDB1 distribution in *w* KD embryo.

Movie S5. GFP-SetDB1 distribution in *Gcn5* KD embryo.

Movie S6. GFP-SetDB1 accumulation at 359-bp foci in *w* KD embryo.

Movie S7. GFP-SetDB1 accumulation at 359-bp foci in *Gcn5* KD embryo.

Movie S8. Control embryo microinjected with DMSO.

Movie S9. The embryo microinjected with (R,R)-59 inhibitor.

Movie S10. GFP-SetDB1^WT^ accumulation at 359-bp foci.

Movie S11. GFP-SetDB1^F578A^ accumulation at 359-bp foci.

## References

Allshire, R.C., and Madhani, H.D. (2018). Ten principles of heterochromatin formation and function. Nat Rev Mol Cell Biol 19, 229–244. 10.1038/nrm.2017.119.

Armstrong, R.L., and Duronio, R.J. (2019). Phasing in heterochromatin during development. Genes Dev 33, 379–381. 10.1101/gad.324731.119.

Batki, J., Schnabl, J., Wang, J., Handler, D., Andreev, V.I., Stieger, C.E., Novatchkova, M., Lampersberger, L., Kauneckaite, K., Xie, W., et al. (2019). The nascent RNA binding complex SFiNX licenses piRNA-guided heterochromatin formation. Nat Struct Mol Biol 26, 720–731. 10.1038/s41594-019-0270-6.

Campos, E.I., Stafford, J.M., and Reinberg, D. (2014). Epigenetic inheritance: histone bookmarks across generations. Trends Cell Biol 24, 664–674. 10.1016/j.tcb.2014.08.004.

Carre, C., Szymczak, D., Pidoux, J., and Antoniewski, C. (2005). The histone H3 acetylase dGcn5 is a key player in Drosophila melanogaster metamorphosis. Molecular and cellular biology 25, 8228–8238. 10.1128/MCB.25.18.8228-8238.2005.

Davis, I.W., Murray, L.W., Richardson, J.S., and Richardson, D.C. (2004). MOLPROBITY: structure validation and all-atom contact analysis for nucleic acids and their complexes. Nucleic Acids Res 32, W615–619. 10.1093/nar/gkh398.

De Rubertis, F., Kadosh, D., Henchoz, S., Pauli, D., Reuter, G., Struhl, K., and Spierer, P. (1996). The histone deacetylase RPD3 counteracts genomic silencing in Drosophila and yeast. Nature 384, 589–591. 10.1038/384589a0.

Dobin, A., Davis, C.A., Schlesinger, F., Drenkow, J., Zaleski, C., Jha, S., Batut, P., Chaisson, M., and Gingeras, T.R. (2013). STAR: ultrafast universal RNA-seq aligner. Bioinformatics 29, 15–21. 10.1093/bioinformatics/bts635.

Elgin, S.C., and Reuter, G. (2013). Position-effect variegation, heterochromatin formation, and gene silencing in Drosophila. Csh Perspect Biol 5, a017780. 10.1101/cshperspect.a017780.

Emsley, P., Lohkamp, B., Scott, W.G., and Cowtan, K. (2010). Features and development of Coot. Acta Crystallogr D Biol Crystallogr 66, 486–501. 10.1107/S0907444910007493.

Fabry, M.H., Falconio, F.A., Joud, F., Lythgoe, E.K., Czech, B., and Hannon, G.J. (2021). Maternally inherited piRNAs direct transient heterochromatin formation at active transposons during early Drosophila embryogenesis. eLife 10. 10.7554/eLife.68573.

Filion, G.J., van Bemmel, J.G., Braunschweig, U., Talhout, W., Kind, J., Ward, L.D., Brugman, W., de Castro, I.J., Kerkhoven, R.M., Bussemaker, H.J., and van Steensel, B. (2010). Systematic protein location mapping reveals five principal chromatin types in Drosophila cells. Cell 143, 212–224. 10.1016/j.cell.2010.09.009.

Fukuda, K., and Shinkai, Y. (2020). SETDB1-Mediated Silencing of Retroelements. Viruses 12. 10.3390/v12060596.

Gaskill, M.M., Soluri, I.V., Branks, A.E., Boka, A.P., Stadler, M.R., Vietor, K., Huang, H.S., Gibson, T.J., Mukherjee, A., Mir, M., et al. (2023). Localization of the Drosophila pioneer factor GAF to subnuclear foci is driven by DNA binding and required to silence satellite repeat expression. Dev Cell 58, 1610–1624 e1618. 10.1016/j.devcel.2023.06.010.

Gu, T., and Elgin, S.C. (2013). Maternal depletion of Piwi, a component of the RNAi system, impacts heterochromatin formation in Drosophila. Plos Genet 9, e1003780. 10.1371/journal.pgen.1003780.

Guo, Y., Mao, X., Xiong, L., Xia, A., You, J., Lin, G., Wu, C., Huang, L., Wang, Y., and Yang, S. (2021). Structure-Guided Discovery of a Potent and Selective Cell-Active Inhibitor of SETDB1 Tudor Domain. Angew Chem Int Edit 60, 8760–8765. 10.1002/anie.202017200.

Jenuwein, T., and Allis, C.D. (2001). Translating the histone code. Science 293, 1074–1080. 10.1126/science.1063127.

Jurkowska, R.Z., Qin, S., Kungulovski, G., Tempel, W., Liu, Y., Bashtrykov, P., Stiefelmaier, J., Jurkowski, T.P., Kudithipudi, S., Weirich, S., et al. (2017). H3K14ac is linked to methylation of H3K9 by the triple Tudor domain of SETDB1. Nat Commun 8, 2057. 10.1038/s41467-017-02259-9.

Kabsch, W. (2010). Xds. Acta Crystallogr D Biol Crystallogr 66, 125–132. 10.1107/S0907444909047337.

Karmodiya, K., Krebs, A.R., Oulad-Abdelghani, M., Kimura, H., and Tora, L. (2012). H3K9 and H3K14 acetylation co-occur at many gene regulatory elements, while H3K14ac marks a subset of inactive inducible promoters in mouse embryonic stem cells. BMC genomics 13, 424. 10.1186/1471-2164-13-424.

Kharchenko, P.V., Alekseyenko, A.A., Schwartz, Y.B., Minoda, A., Riddle, N.C., Ernst, J., Sabo, P.J., Larschan, E., Gorchakov, A.A., Gu, T., et al. (2011). Comprehensive analysis of the chromatin landscape in Drosophila melanogaster. Nature 471, 480–485. 10.1038/nature09725.

Li, X.Y., Harrison, M.M., Villalta, J.E., Kaplan, T., and Eisen, M.B. (2014). Establishment of regions of genomic activity during the Drosophila maternal to zygotic transition. eLife 3. 10.7554/eLife.03737.

Liu, J., Ali, M., and Zhou, Q. (2020). Establishment and evolution of heterochromatin. Annals of the New York Academy of Sciences 1476, 59–77. 10.1111/nyas.14303.

McCoy, A.J., Grosse-Kunstleve, R.W., Adams, P.D., Winn, M.D., Storoni, L.C., and Read, R.J. (2007). Phaser crystallographic software. J Appl Crystallogr 40, 658–674. 10.1107/S0021889807021206.

McKenna, A., Hanna, M., Banks, E., Sivachenko, A., Cibulskis, K., Kernytsky, A., Garimella, K., Altshuler, D., Gabriel, S., Daly, M., and DePristo, M.A. (2010). The Genome Analysis Toolkit: a MapReduce framework for analyzing next-generation DNA sequencing data. Genome Res 20, 1297–1303. 10.1101/gr.107524.110.

Mugat, B., Nicot, S., Varela-Chavez, C., Jourdan, C., Sato, K., Basyuk, E., Juge, F., Siomi, M.C., Pelisson, A., and Chambeyron, S. (2020). The Mi-2 nucleosome remodeler and the Rpd3 histone deacetylase are involved in piRNA-guided heterochromatin formation. Nat Commun 11, 2818. 10.1038/s41467-020-16635-5.

Ninova, M., Chen, Y.A., Godneeva, B., Rogers, A.K., Luo, Y., Fejes Toth, K., and Aravin, A.A. (2020a). Su(var)2-10 and the SUMO Pathway Link piRNA-Guided Target Recognition to Chromatin Silencing. Mol Cell 77, 556–570 e556. 10.1016/j.molcel.2019.11.012.

Ninova, M., Godneeva, B., Chen, Y.A., Luo, Y., Prakash, S.J., Jankovics, F., Erdelyi, M., Aravin, A.A., and Fejes Toth, K. (2020b). The SUMO Ligase Su(var)2-10 Controls Hetero- and Euchromatic Gene Expression via Establishing H3K9 Trimethylation and Negative Feedback Regulation. Mol Cell 77, 571–585 e574. 10.1016/j.molcel.2019.09.033.

Niu, K., Liu, R., and Liu, N. (2018). Quantitative ChIP-seq by Adding Spike-in from Another Species. BIO-PROTOCOL 8.

Orlando, D.A., Chen, M.W., Brown, V.E., Solanki, S., Choi, Y.J., Olson, E.R., Fritz, C.C., Bradner, J.E., and Guenther, M.G. (2014). Quantitative ChIP-Seq normalization reveals global modulation of the epigenome. Cell Rep 9, 1163–1170. 10.1016/j.celrep.2014.10.018.

Otwinowski, Z., and Minor, W. (1997). Processing of X-ray diffraction data collected in oscillation mode. Methods Enzymol 276, 307–326. 10.1016/S0076-6879(97)76066-X.

Padeken, J., Methot, S.P., and Gasser, S.M. (2022). Establishment of H3K9-methylated heterochromatin and its functions in tissue differentiation and maintenance. Nat Rev Mol Cell Biol 23, 623-640. 10.1038/s41580-022-00483-w.

Price, A.J., Manjegowda, M.C., Kain, J., Anandh, S., and Bochkis, I.M. (2020). Hdac3, Setdb1, and Kap1 mark H3K9me3/H3K14ac bivalent regions in young and aged liver. Aging Cell 19, e13092. 10.1111/acel.13092.

Ramirez, F., Dundar, F., Diehl, S., Gruning, B.A., and Manke, T. (2014). deepTools: a flexible platform for exploring deep-sequencing data. Nucleic Acids Res 42, W187–191. 10.1093/nar/gku365.

Regadas, I., Dahlberg, O., Vaid, R., Ho, O., Belikov, S., Dixit, G., Deindl, S., Wen, J., and Mannervik, M. (2021). A unique histone 3 lysine 14 chromatin signature underlies tissue-specific gene regulation. Mol Cell 81, 1766–1780 e1710. 10.1016/j.molcel.2021.01.041.

Samata, M., Alexiadis, A., Richard, G., Georgiev, P., Nuebler, J., Kulkarni, T., Renschler, G., Basilicata, M.F., Zenk, F.L., Shvedunova, M., et al. (2020). Intergenerationally Maintained Histone H4 Lysine 16 Acetylation Is Instructive for Future Gene Activation. Cell 182, 127–144 e123. 10.1016/j.cell.2020.05.026.

Sarot, E., Payen-Groschene, G., Bucheton, A., and Pelisson, A. (2004). Evidence for a piwi-dependent RNA silencing of the gypsy endogenous retrovirus by the Drosophila melanogaster flamenco gene. Genetics 166, 1313–1321. 10.1534/genetics.166.3.1313.

Seller, C.A., Cho, C.Y., and O’Farrell, P.H. (2019). Rapid embryonic cell cycles defer the establishment of heterochromatin by Eggless/SetDB1 in Drosophila. Genes Dev 33, 403–417. 10.1101/gad.321646.118.

Seller, C.A., and O’Farrell, P.H. (2018). Rif1 prolongs the embryonic S phase at the Drosophila mid-blastula transition. PLoS Biol 16, e2005687. 10.1371/journal.pbio.2005687.

Shermoen, A.W., McCleland, M.L., and O’Farrell, P.H. (2010). Developmental control of late replication and S phase length. Curr Biol 20, 2067–2077. 10.1016/j.cub.2010.10.021.

Smart, O.S., Womack, T.O., Flensburg, C., Keller, P., Paciorek, W., Sharff, A., Vonrhein, C., and Bricogne, G. (2012). Exploiting structure similarity in refinement: automated NCS and target-structure restraints in BUSTER. Acta Crystallogr D Biol Crystallogr 68, 368–380. 10.1107/S0907444911056058.

Strom, A.R., Emelyanov, A.V., Mir, M., Fyodorov, D.V., Darzacq, X., and Karpen, G.H. (2017). Phase separation drives heterochromatin domain formation. Nature 547, 241–245. 10.1038/nature22989.

Tang, R., Jiang, Z., Chen, F., Yu, W., Fan, K., Tan, J., Zhang, Z., Liu, X., Li, P., and Yuan, K. (2020). The Kinase Activity of Drosophila BubR1 Is Required for Insulin Signaling-Dependent Stem Cell Maintenance. Cell Rep 31, 107794. 10.1016/j.celrep.2020.107794.

Torres-Zelada, E.F., George, S., Blum, H.R., and Weake, V.M. (2022). Chiffon triggers global histone H3 acetylation and expression of developmental genes in Drosophila embryos. J Cell Sci 135. 10.1242/jcs.259132.

Torres-Zelada, E.F., and Weake, V.M. (2021). The Gcn5 complexes in Drosophila as a model for metazoa. Biochimica et biophysica acta. Gene regulatory mechanisms 1864, 194610. 10.1016/j.bbagrm.2020.194610.

Yuan, K., and O’Farrell, P.H. (2016). TALE-light imaging reveals maternally guided, H3K9me2/3-independent emergence of functional heterochromatin in Drosophila embryos. Genes Dev 30, 579–593. 10.1101/gad.272237.115.

Yuan, K., Seller, C.A., Shermoen, A.W., and O’Farrell, P.H. (2016). Timing the Drosophila Mid-Blastula Transition: A Cell Cycle-Centered View. Trends Genet 32, 496–507. 10.1016/j.tig.2016.05.006.

Zhang, W., Zhang, X., Xue, Z., Li, Y., Ma, Q., Ren, X., Zhang, J., Yang, S., Yang, L., Wu, M., et al. (2019). Probing the Function of Metazoan Histones with a Systematic Library of H3 and H4 Mutants. Dev Cell 48, 406–419 e405. 10.1016/j.devcel.2018.11.047.

Zhao, K., Cheng, S., Miao, N., Xu, P., Lu, X., Zhang, Y., Wang, M., Ouyang, X., Yuan, X., Liu, W., et al. (2019). A Pandas complex adapted for piRNA-guided transcriptional silencing and heterochromatin formation. Nat Cell Biol 21, 1261–1272. 10.1038/s41556-019-0396-0.

